# CAPG: Comprehensive Allopolyploid Genotyper

**DOI:** 10.1101/2022.04.21.488948

**Authors:** Roshan Kulkarni, Yudi Zhang, Steven B. Cannon, Karin S. Dorman

**Affiliations:** Department of Agronomy, Iowa State University, Ames, IA; ORISE Fellow, Corn Insects and Crop Genetics Research Unit, USDA-ARS, Ames, IA; Department of Statistics, Iowa State University, Ames, IA; USDA - Agricultural Research Service, Corn Insects and Crop Genetics Research Unit, Ames, IA; Department of Genetics, Development and Cell Biology, Iowa State University, Ames, IA

## Abstract

**Motivation:** Genotyping by sequencing is a powerful tool for investigating genetic variation in plants, but many economically important plants are allopolyploids, where homoeologous similarity obscures the subgenomic origin of reads and confounds allelic and homoeologous SNPs. Recent polyploid genotyping methods use allelic frequencies, rate of heterozygosity, parental cross or other information to resolve read assignment, but good subgenomic references offer the most direct information. The typical strategy aligns reads to the joint reference, performs diploid genotyping within each subgenome, and filters the results, but persistent read misassignment results in an excess of false heterozygous calls.

**Results:** We introduce the Comprehensive Allopolyploid Genotyper (CAPG), which formulates an explicit likelihood to weight read alignments against both subgenomic references and genotype individual allopolyploids from whole genome resequencing (WGS) data. We demonstrate CAPG in allotetraploids, where it performs better than GATK’s HaplotypeCaller applied to reads aligned to the combined subgenomic references.

**Availability:** Code and tutorials are available at https://github.com/Kkulkarni1/CAPG.git.

## 1 Introduction

Polyploidy is an important phenomenon, especially in plants, that drives the pace and opportunity for evolution in affected lineages (Soltis and Soltis, 2012; Wendel, 2015). The majority of polyploid plants are allopolyploids (rather than autopolyploids), arising due to interspecific hybridization (Parisod *et al*., 2010). Allopolyploids include economically important crops such as peanut, wheat, cotton, quinoa, and rapeseed. While allopolyploidy is common and consequential, available genotyping methods frequently perform poorly in allopolyploid species (Limborg *et al*., 2016; Mason, 2015), causing problems for both applied work (e.g. breeding (Clevenger and Ozias-Akins, 2015)) and basic biology (Kulkarni *et al*., 2020). Figure 1 shows the two classes of SNPs in allopolyploids. A homoeologous SNP is an allelic difference between subgenomes that is not segregating within either subgenome. A homologous or allelic SNP is segregating within at least one subgenome. In allopolyploids, SNPs are often misclassified, typically manifesting as an excess of heterozygous calls (Kulkarni *et al*., 2020; Shirasawa *et al*., 2016).

**Figure 1.**
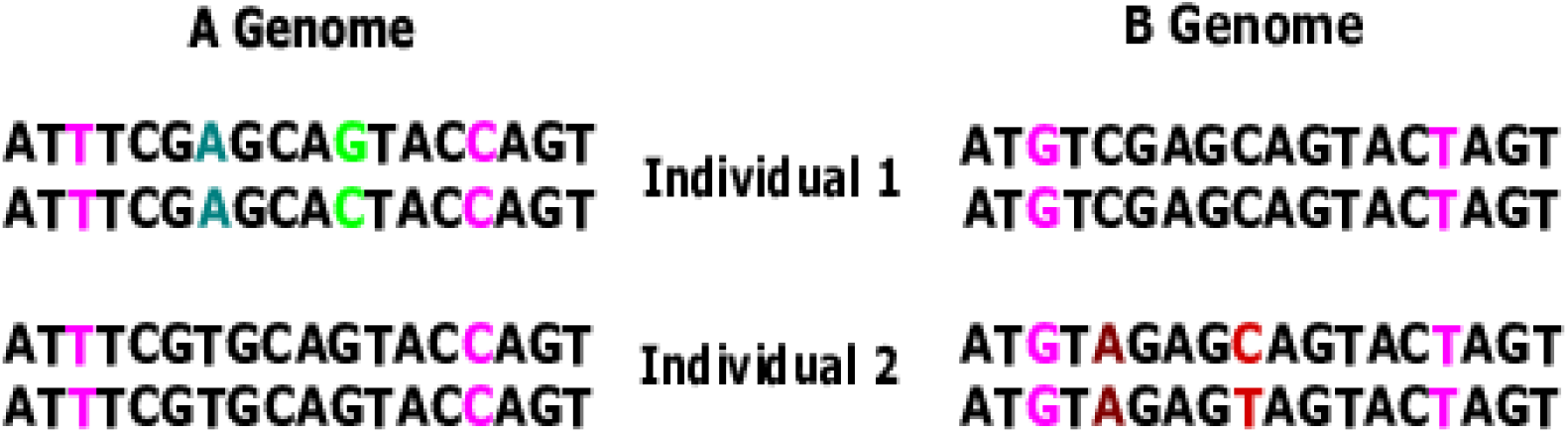
Distinguishing homoeologous and allelic SNPs. Allotetraploid genomes for two individuals, subgenome A (left), subgenome B (right). Pink loci are homoeologous SNPs, different between and constant within subgenomes. Other colored loci are allelic SNPs, green in subgenome A, red in subgenome B. The dark green and brown loci are homozygous. The light green and red loci are heterozygous in one of the individuals.

Allopolyploids can be genotyped given Next Generation Sequencing (NGS) reads aligned to a single reference representing both subgenomes along with allele frequencies, rate of heterozygosity, parental genotype, or other input. The additional input provides information for assigning reads to their subgenomic source and genotyping. For example, EGB (Blischak *et al*., 2018) requires allele frequency estimates from at least one parent species, while updog (Gerard *et al*., 2018) and polyRAD (Clark *et al*., 2019) assume polyploid individuals are sampled from populations with known genetic structure, such as Hardy-Weinberg equilibrium. Without such information, the total alternate allele dosage across both subgenomes can be estimated (Blischak *et al*., 2018) using autopolyploid genotypers, like samtools (Li *et al*., 2009) or Genome Analysis Toolkit (GATK) (McKenna *et al*., 2010). Increasingly, however, available subgenome references (Bertioli *et al*., 2016; Lu *et al*., 2019; Wang *et al*., 2019) or genomes of closely related diploid ancestral species (Bertioli *et al*., 2016; Du *et al*., 2018) provide direct information for assigning reads. SWEEP (Clevenger *et al*., 2015) and HAPLOSWEEP (Clevenger *et al*., 2018), the latter recommended over SWEEP (Peng *et al*., 2020), use these references to identify allelic SNPs via homozygous individuals at variable sites bracketed by likely homoeologous loci. However, the most common subgenome reference aware method (Peng *et al*., 2017; Zhou *et al*., 2014, and M2 in Peng *et al*., 2020) aligns reads to both subgenomes, keeps uniquely aligned reads, and applies a diploid genotyper to each subgenome.

Methods using subgenomic references either use an aligner to imperfectly partition reads to subgenomes or incorrectly process reads from all allopolyploid chromosomes with a diploid genotyper. Instead, we describe the Comprehensive Allopolyploid Genotyper (CAPG), which uses a likelihood to weight read alignments to *both* subgenomes while genotyping individuals from NGS data. Calls are reported in Variant Call format (VCF), with familiar measures, such as genotype likelihood, to gauge statistical support. For samples of individuals, loci are classified as homoeologous SNPs, allelic SNPs within subgenome, or invariant. We test the method on simulated data and whole-genome sequencing (WGS) data from two allotetraploids: Peanut (*Arachis hypogaea*) and Cotton (*Gossypium hirsutum*), comparing to GATK on reads partitioned by joint alignment to subgenomic references. HAPLOSWEEP could not run because no haplotypes passed the inclusion criteria. While currently implemented for allotetraploids, CAPG can be extended to higher ploidy levels.

## 2 Approach

### 2.1 Model

Consider a homoeologous genomic region in an allotetraploid individual with A and B subgenomes. We assume reference sequences, with known alignment in the homoeologous region, are available for both subgenomes. Our goal is to genotype the individual in the homoeologous region given whole genome sequencing reads. We align each read, once each to subgenome A and B, producing *n* reads ***r***_1_, ***r***_2_, …, ***r***_*n*_ and quality scores ***q***_1_, ***q***_2_, …, ***q***_*n*_ with *homoeologous alignments* to a locus in the homoeologous region. Two alignments are homoeologous at a locus if the *same* read base aligns to the homoeologous locus in both subgenomes. All other read alignments spanning this locus are discarded as likely sequencing or library preparation errors, such as recombination, or reads of paralogous regions, any such reason rendering contradictory alignments. Discarded reads may also reflect genuine alignment ambiguity, particularly around indels, where alignment refinement is warranted (McKenna *et al*., 2010). CAPG does not genotype indels.

The genotype we wish to call, for example CC/CT, represents the unordered nucleotides at a locus from the maternal and paternal copies of the A subgenome, followed by the unordered nucleotides of the B subgenome. Assuming no more than two distinct nucleotides at the locus (for want of better terminology, the major and minor alleles), we can represent the genotype as ***M*** = (*m*_1_, *m*_2_), where *m*_1_, *m*_2_ *∈* {0, 1, 2} are the numbers of minor alleles in the A and B subgenomes. Given *n* independent reads with homoeologous alignments to a locus, we seek the genotype ***M*** with highest posterior probability,

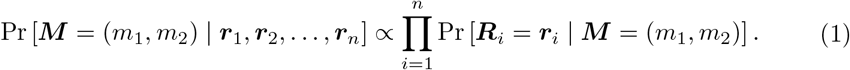

Conditioning on the homoeologous alignments of the *i*th read and assuming only the true source subgenome *S*_*i*_ *∈* {1, 2*}, i.e*., which is the true alignment, is unknown, the read likelihood is

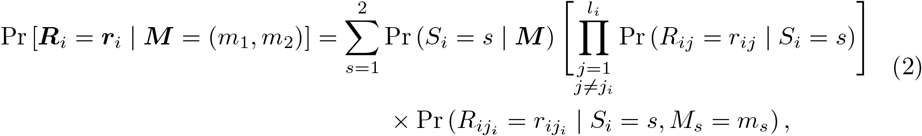

where *j*_*i*_ is the site in read ***r***_*i*_ aligned to the locus in both alignments and *l*_*i*_ is the aligned length of the *i*th read after removing all insertions, deletions and sites without homoeologous alignments. While read indels are highly informative of subgenomic source because sequencing indels are rare, we neglect them out of concern that unrecognized homoeologous indel variation not present in the references or segregating homologous indel variation would drive the read likelihood. Thus, we solely rely on homoeologous (mis)matches to provide the signal for the subgenomic assignment of reads. When homoeologous SNPs are sparse, the genotype will be called with appropriate uncertainty.

We finish by formulating each probability in the equation. Assuming uniform subgenomic coverage, Pr (*S*_*i*_ = *s* | ***M***) = 0.5, let *t*_*r*_ be the major and *t*_*a*_ the minor allele at the locus. If *T*_*i*_ is the true nucleotide in the chromosome sequenced in the *i*th read, then under the additional assumption of equal chromosomal coverage,

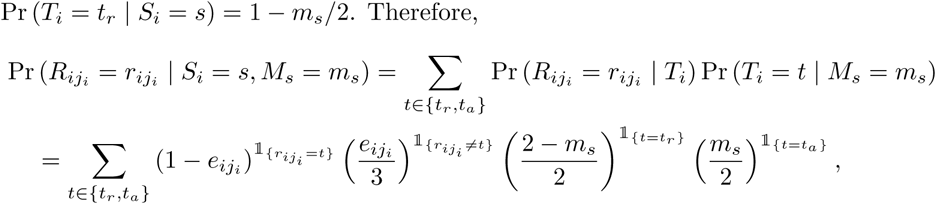

where 𝟙_{·}_ is an indicator function, 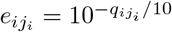 is the probability of a sequencing error assuming Phred quality scores (Ewing and Green, 1998) and assuming equal probability of the three possible substitution errors. For read position *j* ≠ *j*_*i*_ in read *i* aligned to locus *l*, we assume the genotype is homozygous *g*_*sl*_*g*_*sl*_, where *g*_*sl*_ is the allele in the *s*th reference genome at locus *l*. Thus, Pr(*R*_*ij*_ = *r*_*ij*_ | *S*_*i*_ = *s*) is

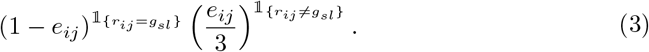

The assumption of homozygosity at other loci may be violated by nearby allelic SNPs or errors in the references, and Phred errors are not warranted in most datasets. Reference correction and quality score recalibration can help (McKenna *et al*., 2010), but we leave such efforts to future work.

### 2.2 Genotyping and SNP calling

To genotype an individual is to identify the most plausible alleles at each genomic locus. SNP calling considers data from multiple individuals to identify variable loci in the population. Modern SNP callers typically assume and estimate a probability distribution, *e.g*., Hardy-Weinberg equilibrium (HWE), for genotypes at a locus in a population. Estimating population parameters from the data of multiple individuals typically improves genotype calling, especially in low coverage situations (Nielsen *et al*., 2011). Such an approach is possible in our framework, but for our data, we could not assume Hardy-Weinberg equilibrium, let alone a common source population. Instead, we independently genotype individuals and combine the results to perform SNP calling assuming a uniform (uninformative) prior on the genotypes. We leave it to future work to model and estimate parameters of the population.

#### 2.2.1 Genotyping

Sampled individuals are independently genotyped assuming biallelic loci. For each locus in a homoeologous region, we identify the two most common alleles observed among the *n*_*k*_ reads 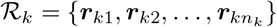 with homoeologous alignments to the locus in individual *k*, calling the most common allele the *major allele t*_*kr*_ and the second most common allele the *minor allele t*_*ka*_, breaking ties by alphabetic ordering of the nucleotides. If there is no second allele observed in the reads with homoeologous alignments, we choose the first alphabetically ordered nucleotide not already denoted the major allele as the minor allele. We then compute the posterior probability of all nine possible allotetraploid genotypes via Eq. (1) and call the genotype as the most likely. We assess the support for heterozygosity at the locus in individual *k* and subgenome *g* as the log likelihood ratio

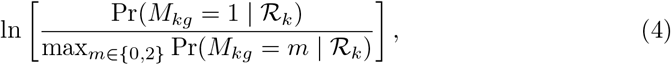

where *M*_*kg*_ is the genotype for individual *k* at the locus in subgenome *g*.

#### 2.2.2 SNP calling

We also limit SNP identification to biallelic SNPs, involving nucleotides *N*_1_ and *N*_2_. We compute metrics to call homoeologous and allelic SNPs under a uniform prior over all possible genotypes. Since we identify major *t*_*kr*_ and minor *t*_*ka*_ alleles separately for each individual *k*, if the posterior probability of the required genotype is not among the nine computed, we substitute the minimum of the nine. This approximation can be remedied by defining the major and minor alleles from the joint data.

The support for allelic SNPs in subgenome *g* is assessed as

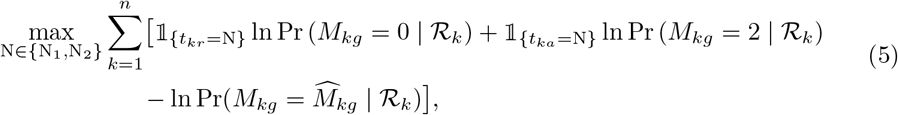

where 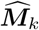 is the called genotype for individual *k* at the locus. The sum allows either major or minor allele to be N, which varies with read coverage. Support for homoeologous SNPs is assessed as

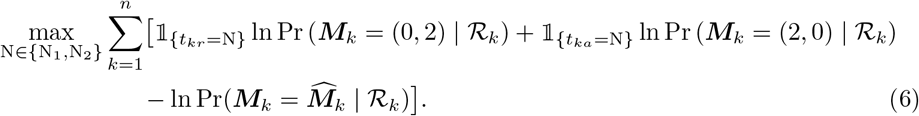

## 3 Methods

### 3.1 Implementation

Figure 2 describes the main ideas behind CAPG genotyping. Reads are aligned separately to both subgenomic reference genomes, major and minor alleles for a locus are identified, and the likelihood of each read aligned to each subgenome for nine possible genotypes is used to compute the posterior probabilities by Eq. (1). There is opportunity to modify the workflow, from choice of short read aligner to read filtering or skip genotyping loci with, for example, low coverage (details in S1.1). Worked examples are available at https://github.com/Kkulkarni1/CAPG. In this work, we genotype all loci except indels in the subgenome reference alignment and those with no homoeologous coverage of one or more subgenomes by Eq. (S1).

**Figure 2.**
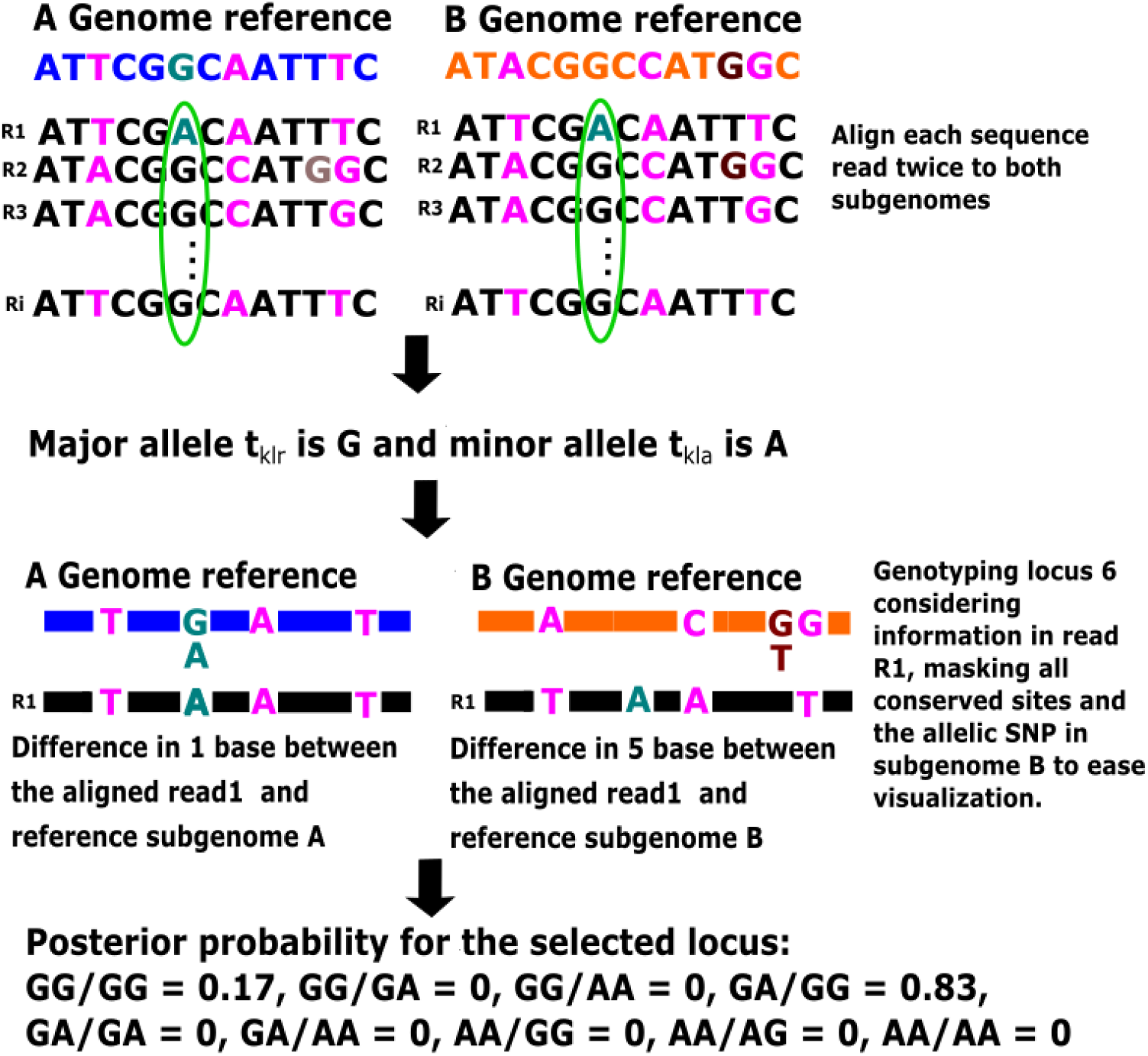
Genotyping an allelic SNP with CAPG. Blue: subgenome A reference sequence; Orange: subgenome B reference sequence; Pink: homoeologous SNPs at loci 3, 8, and 12; Green: segregating allele in subgenome A at locus 6; Brown: segregating allele in subgenome B at locus 11; Green ellipse identifies the locus to genotype in this example, with genotype GA/GG yielding the highest posterior probability of 0.83.

CAPG includes optional post-hoc filters for allotetraploid genotyping in the face of data artefacts. The model (§2.1) assumes equal homoeologous and homologous chromosome coverage, so a false heterozygous call may arise at a locus with an unusually high sequencing error rate, an amplified PCR error, or biased sequencing of one subgenome. Though the resulting high frequency of the alternate allele is inconsistent with variation due to sequencing errors, it is unlikely to exactly match 50% of the subgenomic coverage. While the assumption of equal chromosomal coverage is not always valid (Gerard *et al*., 2018; Rozowsky *et al*., 2011), we derive and implement a likelihood ratio test (details in §S1.1.2) of equal homologous coverage in the presence of unequal homoeologous coverage, which can be used to screen heterozygous calls for some error and coverage artefacts.

### 3.2 Simulation

We compared CAPG and GATK (McKenna *et al*., 2010) on their ability to provide valid input, *e.g*., normalized PHRED-scaled likelihood (PL), for correctly identifying heterozygous loci, allelic SNPs, and homoeologous SNPs on simulated WGS data. Simulation details are in Supplement §S1.3. Briefly, we simulated 100,000 loci in 50 individuals, with about 1 allelic SNP per 100 loci and homoeologous SNP rate *r*_*h*_ *∈* {0.005, 0.007, 0.01}. We genotyped each locus in each individual based on simulated paired end reads of length 150 from fragments of mean length 300 *±* 10 and subgenomic coverage rate *c* ∈ {10, 20, 40}.

#### 3.2.1 Running CAPG

Simulated reads were aligned separately against the simulated reference subenomes using BWA-MEM2 (Vasimuddin *et al*., 2019) with default settings. Some loci (less than 0.1%) were dropped from heterozygosity calling because of no read coverage; two loci were dropped from SNP calling because of no read coverage across all 50 individuals at coverage level *c* = 5 (details in Table S1). These loci are excluded from all presented results. PR curves were plotted based on the CAPG metrics of Eqs. (4)–(6) for evaluating heterozygosity, allelic SNPs, and homoeologous SNPs. The metric (6) is sometimes −∞, which is replaced in plots with 2*r*_(2)_ − *r*_(3)_, where *r*_(*i*)_ is the *i*th smallest metric value. Loci are sometimes subsampled as indicated in figure legends to ease viewing.

#### 3.2.2 Running GATK

We aligned reads to the joint tetraploid reference A *and* B subgenomes, using BWA-MEM2, followed by discard of secondary alignments. GATK’s HaplotypeCaller was used to separately genotype each subgenome using default parameters. Quantities equivalent to CAPG metrics Eqs. (4)–(6) (see Eqs. (S3)–(S5)) were computed for all homoeologous positions in the subgenomic reference alignment.

### 3.3 Validation on real data

We downloaded WGS resequencing data from 14 peanut and 9 cotton germplasms of diverse origin (Clevenger *et al*., 2017; Fang *et al*., 2017; Pan *et al*., 2020) (accessions listed in Tables S3 and S4). We identified genic regions in the genome annotation files of the peanut Tiffrunner assembly (Bertioli *et al*., 2019) and the cotton TM1 assembly (Li *et al*., 2015), downloaded from PeanutBase (Dash *et al*., 2016) and NCBI, respectively. We selected 1,000 genic homoeologies found using BLAST (Altschul *et al*., 1990) with 95–98% sequence similarity and alignment length over 1000 bps. These sequences were genotyped using CAPG as for simulation, but post-hoc filtered for calls with expected coverage of either subgenome below eight (Minimum coverage in §S1.1.1), no subgenome mismatches within read distance (Identifiable in §S1.1.2), or for heterozygous calls, rejection of the equal homologous coverage hypothesis at significance level 0.05 (Equal homologous coverage test in §S1.1.2). GATK was run as for simulation data, except indel variants were removed by bcftools, calls with fewer than eight reads per either subgenome or no subgenomic mismatches within read distance, and heterozygous calls rejected at significance level 0.05 via a likelihood ratio test of equal homologous coverage using the allele depth data (SAM tag AD) reported by HaplotypeCaller were discarded.

## 4 Experimental results

### 4.1 Simulation

To verify CAPG performance, we simulated allotetraploid data while varying subgenomic read coverage (*c*) and homoeologous rate (*r*_*h*_). As expected, performance of CAPG improves with higher coverage and more homoeologous SNPs. The details are provided in the Supplement §S2.1.

We also compared the performance of CAPG with benchmark GATK (McKenna *et al*., 2010), using reads assigned to subgenome by alignment, on ability to detect heterozygosity and predict SNPs. Performance as a function of coverage and homoeologous rate is detailed in §S2.1. Here we report results for the simulation with *r*_*h*_ = 0.7% and *c* = 10. Fig. 3a shows that CAPG better detects heterozygosity than GATK. For allelic SNPs (Fig. 3b), CAPG is only superior to GATK at high precision. At the threshold where the PR curves cross, there are 52 allelic SNPs in subgenome A not called by CAPG and all are nonidentifiable, having no homoeologous SNP within read length distance. The CAPG metric is appropriately low and provides equal support for an allelic SNP in subgenome B. The GATK metric also reflects ambiguity, with about half (30 vs. 22) showing stronger support for an allelic SNP in subgenome B, where the locus is invariant. The difference is that the GATK metrics are larger; 30 of the 52 unidentifiable allelic SNPs are already called at the threshold where the PR curves cross. This strong signal is an artefact of the complete confidence placed in the read assignments, which also results in high rates of false heterozygous calls (Fig. 3a). After removing all nonidentifiable loci, both PR curves improve (solid lines in Fig. 3), but CAPG is superior, only misclassifying two identifiable allelic SNPs using a threshold of 21.1. Finally, CAPG homoeologous SNP calling at the default threshold reaches equal precision but lower recall than GATK (Fig. 3c). The increased uncertainty of CAPG is because it does not use the genotyped locus to assign reads, a fact we discuss further in the next section.

**Figure 3.**
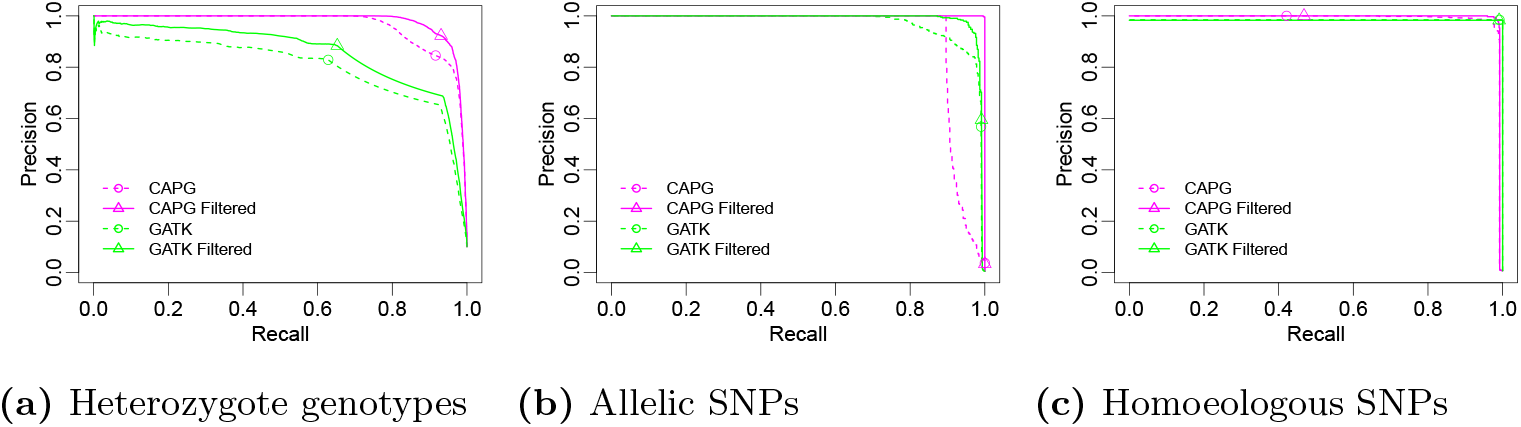
PR curves for CAPG and GATK on simulated data. Performance of CAPG and GATK metrics to identify (a) heterozygous loci in subgenome A, (b) allelic SNPs in subgenome A, and (c) homoeologous SNPs when coverage *c* = 10 and homoeologous rate *r*_*h*_ = 0.007. Heterozygous data are subsampled with all true positive loci and 100,000 randomly sampled true negative loci. Circles/triangles represent the threshold value (0), a liberal choice (high recall, low precision, also see Fig. S4) for genotyping heterozygotes and progressively more conservative with sample size for SNP calling.

A scatter plot of the CAPG and GATK metrics underlying the PR curves is shown in Fig. 4, with true status indicated in color. Nonidentifiable loci are plotted as circles; these loci form the lower tier of points in the homoeologous facet. CAPG metrics are much closer to linearly separable than GATK metrics. GATK sometimes indicates strong support for false positive SNP calls, and while CAPG is not perfect, when it makes a false call, the metric indicates borderline support.

**Figure 4.**
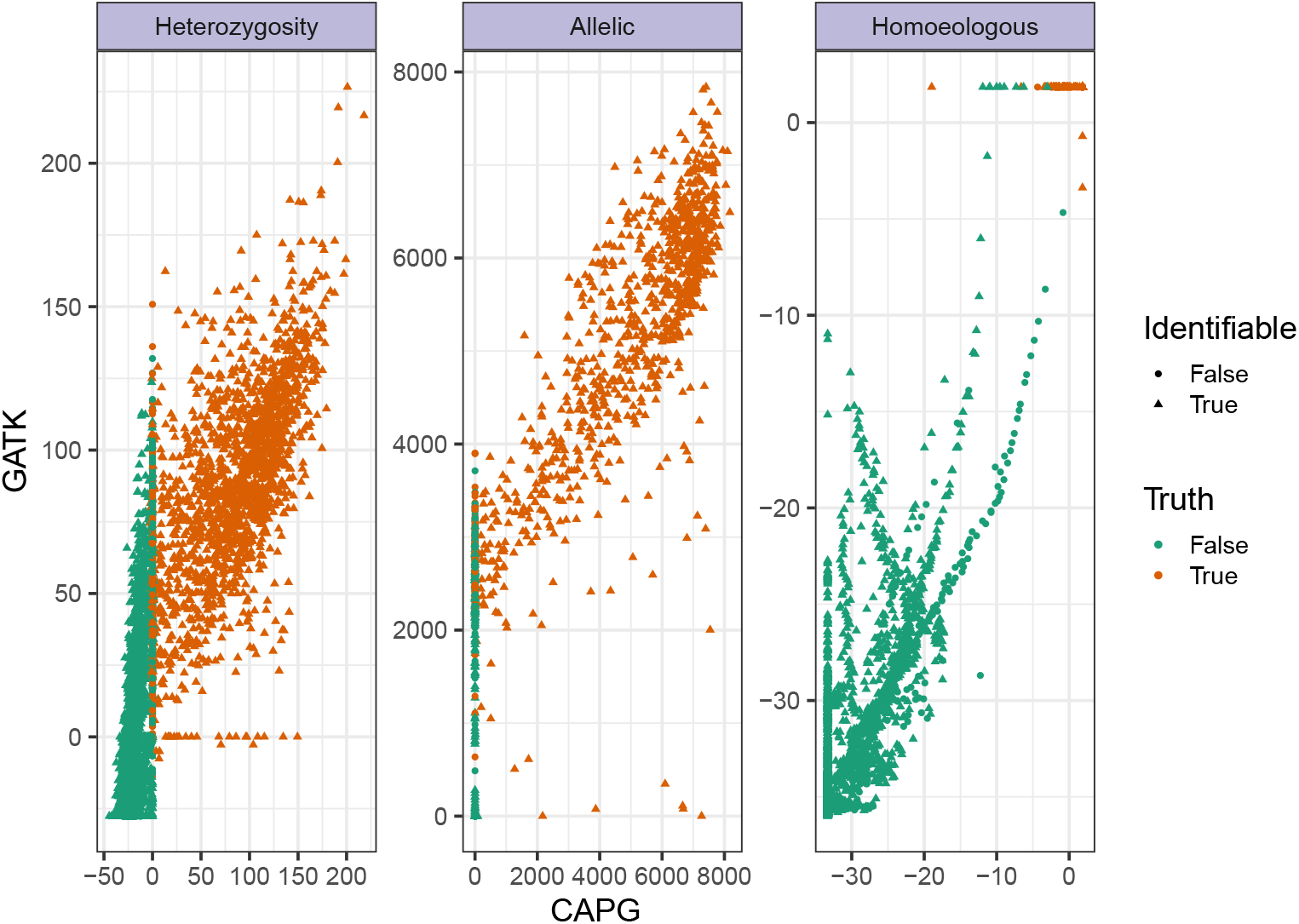
Comparing CAPG and GATK metrics in simulation. Scatter plot of heterozygosity, allelic SNP and homoeologous SNP metrics on simulated data with coverage *c* = 40 and homoeologous rate *r*_*h*_ = 0.007. Homoeologous metrics (Eqs. (6) and (S5)) are transformed via Box-Cox transformation − 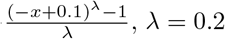 to avoid overplotting at upper right, and the stack of points on the left represent a transformation of CAPG metric value −*∞* (see methods). There remain overplotted true homoeologous SNPs, but all true negatives (green) are visible after transformation. In addition to subsampling done for heterozygosity (see Fig. 3), we further subsample to avoid excess overplotting, keeping all points with CAPG metric *>* 0 for heterozygosity or finite for homoeologous SNPs and subsampling 10% of all other points.

### 4.2 Real data

We collected 14 peanut and 9 cotton accessions. After whole genome alignment, average read coverage per locus per subgenome was 25 (range 15–40 across accessions) for peanut and 12 (range 9–21) for cotton. We genotyped 1,000 selected gene sequences with both CAPG and GATK. Selection of thresholds for calling heterozygous genotypes, allelic SNPs, and homoeologous SNPs is described in §S2.2.

SNPs were distributed throughout the selected genic sequences in both subgenomes of peanut and cotton (data not shown). Table 1 shows 1.7% of genotyped loci in peanut are likely homoeologous SNPs, far fewer are likely allelic SNPs (0.016%, distributed equally in both subgenomes), and even fewer loci are convincingly heterozygous, suggesting the sample consists of largely inbred individuals. We found a higher homoeologous (2.5%) and allelic (0.08%) SNP rate in cotton, consistent with previous findings (Bertioli *et al*., 2016; Fang *et al*., 2017). The chosen threshold probably excludes some true allelic SNPs. While metrics for allelic SNPs and monomorphic loci are well-separated in simulation (Figs. S4h and S4k), the two groups clearly overlap in real data (Figs. S4b and S4e). Nevertheless, there are metrics in both species indicative of allelic SNPs. Higher coverage, larger samples, or biological verification may confirm the predicted allelic SNPs from this study.

**Table 1.** SNPs identified by CAPG from 1000 selected gene sequences among 14 Peanut accessions. Chrom.: genes from listed A chromosome, mostly aligned to homoeologous B chrom.; Total bps: total loci in selected genes; Typed bps: genotyped loci; A: subgenome A; B: subgenome B; H: homoeologous SNP calls; Het.: heterozygous.

| Chrom.<br>A/B | No. of<br>Genes | Total<br>bps | Typed<br>bps | SNP Type |  |  | Het. Loci |  |
| --- | --- | --- | --- | --- | --- | --- | --- | --- |
|  |  |  |  | A | B | H | A | B |
| Chr1/11 | 124 | 382,667 | 374,483 | 27 | 41 | 6,451 | 10 | 28 |
| Chr2/12 | 59 | 191,031 | 187,186 | 10 | 10 | 3,474 | 0 | 1 |
| Chr3/13 | 163 | 504,002 | 495,213 | 38 | 36 | 8,381 | 13 | 11 |
| Chr4/14 | 77 | 248,104 | 240,961 | 48 | 55 | 4,030 | 30 | 20 |
| Chr5/15 | 129 | 388,839 | 380,297 | 18 | 28 | 6,276 | 0 | 8 |
| Chr6/16 | 103 | 335,647 | 329,849 | 19 | 26 | 5,434 | 0 | 13 |
| Chr7/17 | 60 | 195,670 | 190,549 | 26 | 10 | 3,578 | 1 | 1 |
| Chr8/18 | 94 | 284,517 | 273,462 | 21 | 10 | 4,580 | 3 | 0 |
| Chr9/19 | 113 | 380,543 | 369,665 | 32 | 17 | 6,378 | 9 | 1 |
| Chr10/20 | 78 | 265,632 | 259,947 | 16 | 13 | 4,458 | 5 | 2 |
| **Total** | 1,000 | 3,176,652 | 3,101,612 | 255 | 246 | 53,040 | 71 | 85 |

**Table 2.** SNPs identified by CAPG from 1000 selected gene sequences among 9 Cotton accessions. Chrom.: genes from listed subgenome A chromosome, mostly aligned to homoeologous D chrom.; Total bps: total loci in selected genes; Typed bps: genotyped loci; A: subgenome A; D: subgenome D; H: homoeologous SNP calls; Het.: heterozygous.

| Chrom.<br>A/B | No. of<br>Genes | Total<br>bps | Typed<br>bps | SNP Type |  |  | Het. Loci |  |
| --- | --- | --- | --- | --- | --- | --- | --- | --- |
|  |  |  |  | A | B | H | A | B |
| Chr1/14 | 74 | 188,535 | 170,513 | 79 | 94 | 4,517 | 9 | 1 |
| Chr2/15 | 47 | 129,470 | 101,284 | 42 | 34 | 2,966 | 1 | 3 |
| Chr3/16 | 57 | 144,060 | 127,882 | 30 | 58 | 3,188 | 2 | 2 |
| Chr4/17 | 34 | 88,886 | 70,335 | 38 | 45 | 2,142 | 20 | 36 |
| Chr5/18 | 164 | 470,043 | 403,721 | 167 | 197 | 10,591 | 11 | 9 |
| Chr6/19 | 62 | 160,249 | 150,071 | 40 | 39 | 3,680 | 1 | 2 |
| Chr7/20 | 73 | 186,525 | 173,147 | 56 | 125 | 4,283 | 8 | 2 |
| Chr8/21 | 80 | 198,353 | 184,720 | 73 | 61 | 4,624 | 8 | 3 |
| Chr9/22 | 72 | 197,435 | 191,502 | 80 | 66 | 4,497 | 8 | 28 |
| Chr10/23 | 74 | 186,009 | 152,265 | 79 | 59 | 4,396 | 9 | 2 |
| Chr11/24 | 108 | 314,290 | 289,887 | 69 | 67 | 6,880 | 1 | 2 |
| Chr12/25 | 95 | 231,822 | 220,009 | 72 | 96 | 5,255 | 6 | 0 |
| Chr13/26 | 60 | 162,958 | 143,942 | 94 | 46 | 3,640 | 14 | 2 |
| **Total** | 1,000 | 2,658,635 | 2,379,293 | 919 | 987 | 60,659 | 98 | 92 |

Scatterplots of unfiltered CAPG and GATK metrics for the peanut data show greater variability than simulation data (Fig. 5). There is positive association between CAPG and GATK in all three metrics, but the association is weak (Spearman correlation *ρ <* 0.4) for the SNP metrics (Table S5). We sampled several identifiable loci and manually genotyped the read alignments (plot symbols). As for simulated data, there is a pattern of loci with CAPG heterozygosity metric near 0 and increasing GATK metric up the *y*-axis, likely false heterozygous calls by GATK and confirmed in visual spot checks (red crosses). However, we also could not confirm heterozygosity for loci where both methods agree (upper right), mostly because paralogous reads mapping to the locus obscured the signal (gray stars). We could manually identify and ignore paralogous reads at two loci to find no heterozygosity (red crosses), but neither method can automatically remove paralogous reads. CAPG and GATK strongly disagree on a proportion of loci in the upper left (0.4%) and lower right (3%) of the allelic SNP plot. Causes for loci in the lower right include aligned paralogous reads, mismatches between the subgenomes not recapitulated in the reads, or nearly nonidentifiable loci. The latter two kinds of loci also often appear in the upper swath of the homoeologous metric plot, where GATK is certain but CAPG is uncertain in the homoeologous SNP call. These loci tend to be subgenomic mismatches (purple points), information used to separate the reads passed to GATK but ignored by CAPG for read assignment. In our view, CAPG metrics reflect appropriate ambiguity in these cases. In the upper left, GATK is mislead by mismapped reads, low levels of an alternate allele with uncertain provenance, and supplementary alignments. Our filters remove many (62% upper left, 92% lower right) of these cases (Figure S6).

**Figure 5.**
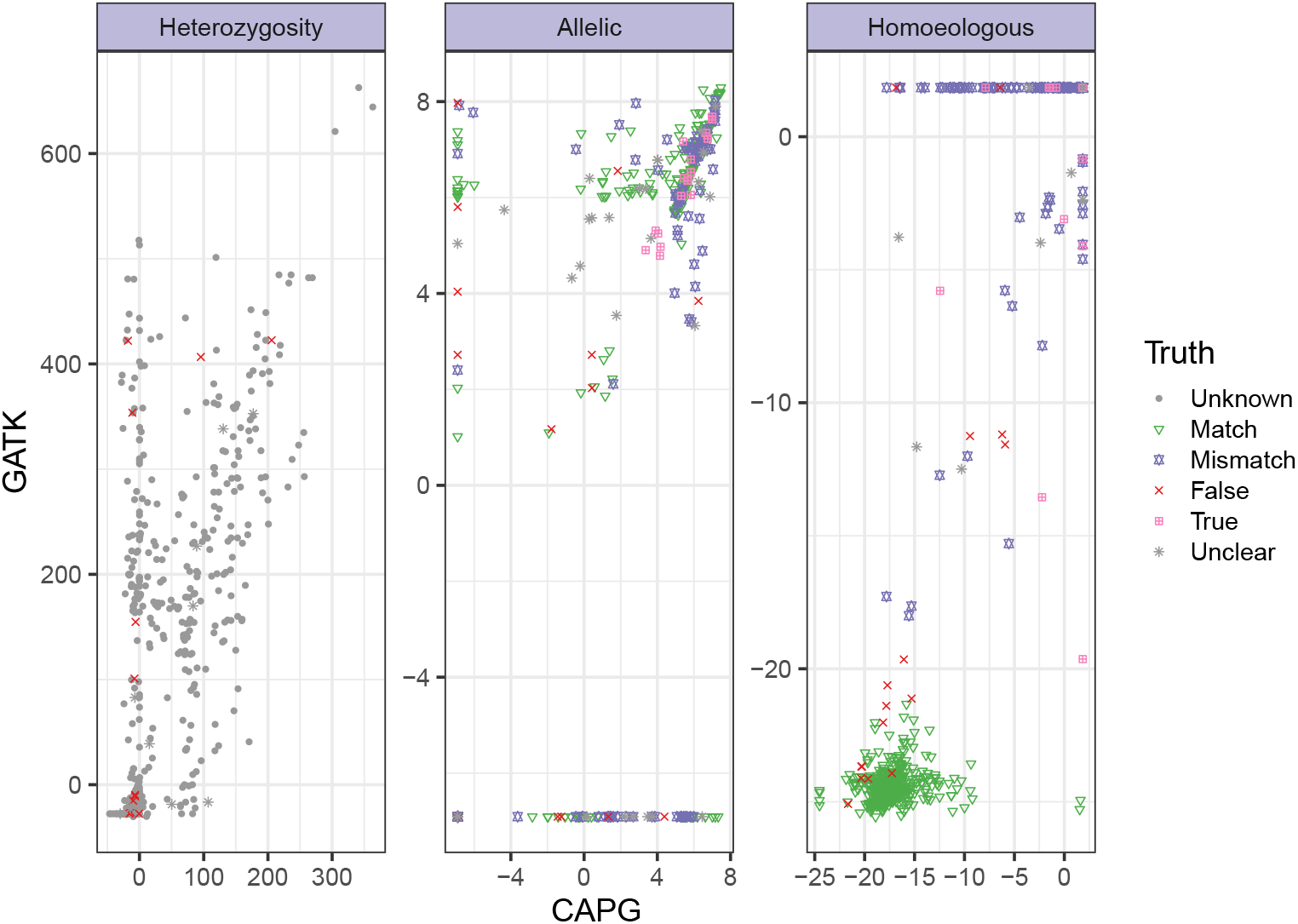
Comparing CAPG and GATK metrics in real peanut data. Scatter plot of metrics for heterozygosity, allelic and homoeologous SNPs. We examine alignments to confirm (True: pink boxed ‘x’), reject (False: red plus) or fail to resolve (Unclear: gray star) a small selection of loci. Otherwise, the heterozygosity status is unknown (gray circle), but we indicate if there is a subgenomic reference nucleotide match (green triangle) or mismatch (purple square) in the allelic and homoeologous facets. After including all hand-verified loci, a stratified sample was taken to over-sample likely heterozygous calls by either method, so 50% of sampled loci have CAPG or GATK metrics above the 99.5th percentile. For allelic SNP metrics, we sampled 25% loci with subgenomic mismatch, 25% loci with either CAPG or GATK metric above the 99.5th percentile, 25% loci with subgenomic match, and 25% with both metrics below the 99.5th percentile. For homeologous SNP metrics we sampled 50% loci with subgenomic mismatch and 50% with subgenomic match and low metrics by CAPG and GATK. For an unbiased view of the metrics, see Figures S7–S9.

We find evidence that CAPG metrics are more tightly associated with other evidence of true SNPs. CAPG metrics better correlate with manually assessed truths (Table S5), but while the sampling strategy for loci to assess was neutral, it was not balanced, so we also examined associations in unsampled data. Mismatches between the reference subgenomes should occur at all homoeologous SNPs and some allelic SNPs, so a mismatch can serve as a noisy label for either SNP. PR curves (Figure S10) and numeric associations (Table S5) demonstrate the CAPG metric better discriminates mismatch-identified allelic SNPs but not homoeologous SNPs. GATK’s apparent superior performance on homoeologous SNPs is an artefact of using the locus itself to partition reads. Examination of allelic and homoeologous SNP metrics in Figures S8 and S9 suggests some subgenomic mismatches are actually allelic SNPs. Finally, we expect more extreme (definitive) metrics at identifiable loci. We use Levene’s statistic to measure spread in the metric, confirming the heterozygous metric (CAPG 5 *×* 10^4^ vs. GATK 1 *×* 10^2^), the allelic metric (CAPG 0.05 vs. GATK 0.01), but not the homoeologous metric (CAPG 73 vs. GATK 85) are more extreme at identifiable loci in CAPG. A full discussion of these results and more is in §S2.2.1.

## 5 Discussion

We propose a likelihood-based genotyper, CAPG, to accurately call genotypes and SNPs in allotetraploids. We have shown that CAPG is better at identifying SNPs in both simulation and real data than the benchmark GATK applied to reads split by alignment to reference subgenomes. We now discuss the advantages and limitations of CAPG.

### 5.1 Likelihood-based genotyping

Li (2011) has shown the value of likelihood methods for genotyping and calling SNPs, and it is logical to extend such models to genotype allopolyploids and account for the unknown subgenomic source of the read. To apply the model, we condition on independent alignments of each read against both reference subgenomes. We use the likelihood to assess their relative support, avoiding the approximate choice between homoeologs made by short read aligners. A complication is that CAPG obtains joint estimates of the allotetraploid genotype, not independent estimates of the two homoeologous diploid genotypes, which necessitates a moderate amount of post-processing to merge into traditional genotyping pipelines. To encourage genotyping at the allotetraploid level, we provide metric scores for allelic and homoeologous SNP calling across the sample. We also provide a formal test of equal coverage of homologous chromosomes that can be used to screen heterozygous calls.

Li (2011) and others have modeled the genotype proportions in the population during variant discovery, and CAPG can be extended to consider such hierarchical models. Our demonstration genotypes individuals independently and then combines the genotyping metrics post-hoc, essentially placing a uniform prior on plausible genotypes (Nielsen *et al*., 2011). This approach is reasonable for our real data examples since they include individuals with unknown relationships, where it is unclear what assumptions to impose about the population.

Our presentation and software focus on allotetraploids, but extension to higher ploidies is straightforward. Ambiguous genotypes are increasingly probable at high ploidy. An allelic SNP linked to the alternate allele at homoeologous SNP (0, 2, 2) is either in subgenome B or C, and CAPG would appropriately communicate the uncertainty.

### 5.2 Model limitations

As expected, the performance of our method declines with decreasing coverage. When appropriate, inclusion of a hierarchical population model can increase power, but with advances in NGS technologies, it should also be possible to obtain sufficient coverage.

All methods require homoeologous sites to assign reads to subgenomes, so accuracy declines with short reads, low homoeologous rate or poor subgenomic references. Competitive alignment against the joint reference can recapitulate support using GATK (not CAPG) for homoeologous loci already represented in the subgenomic references. Generally, GATK predisposes homoeologous SNPs at all subgenomic mismatches, even when they are actually allelic SNPs or errors.

In repetitive genomes, reads from paralogous sources lead to excess evidence of heterozygosity. Post hoc tests of equal homologous coverage in heterozygotes can eliminate some such false calls. It is also possible to check for evidence of more than four haplotypes at a locus, for example using denoisers (Peng and Dorman, 2020) to identify and remove paralogous reads or entire contaminated regions. Such an approach was successful when applying CAPG to amplicon sequences (data not shown).

Both methods condition on alignments of reads to the subgenomic references and of subgenomic references to each other. Incorrect subgenomic read assignment disrupts genotyping by GATK, but both methods may detect false signal around indels when there are multiple plausible alignments. It is possible to refine read alignments prior to genotyping, as has shown promise (McKenna *et al*., 2010). Incorrect subgenomic reference alignments would align non-homoeologous read positions, inducing errors in allelic and homoeologous SNP calling. It is worth investing in good references and alignments. Model modifications could account for known reference deficiencies and computationally intense methods could integrate over uncertainty in the alignment.

### 5.3 Biological significance

Accurate prediction of heterozygosity, allelic SNPs and homoeologous SNPs is important for basic biology – for example, in evolutionary studies – and the applied science of plant breeding and other applications. Since only allelic SNPs segregate, it is clearly important to be able to distinguish allelic and homoeologous SNPs, but accurate genotyping also improves our ability to study gene gain or loss after polyploidization, major structural rearrangements or conservation between homoeologous chromosomes, and functional divergence of polyploids from diploids. From a crop improvement viewpoint, identifying functionally conserved homoeologs can help elucidate the genetic basis for traits of interest.

## Supporting information

Supplementary methods and results.

## Acknowledgments

Thanks to J. F. Wendel for helpful discussion about allotetraploid cotton.

