## Supplementary methods and results. for "CAPG: Comprehensive Allopolyploid Genotyper"

### Supplementary Material for
### "CAPG: Comprehensive Allopolyploid Genotyper"

Roshan Kulkarni[1], Yudi Zhang[2], Steven B. Cannon[3] and Karin S. Dorman[2,4]
[1] *Department of Agronomy, Iowa State University*
[2] *Department of Statistics, Iowa State University*
[3] *USDA - Agricultural Research Service, Corn Insects and Crop Genetics Research Unit*
[4] *Department of Genetics, Development and Cell Biology, Iowa State University*
Ames, IA 50011, USA

### S1 Methods

#### S1.1 Implementation

Section 3.1 and Figure 2 of the main text provide a procedural overview of CAPG genotyping. Here we provide some additional details. CAPG genotypes one homoeologous region at a time. It currently does not index SAM alignment files, so although it can take and process large SAM files, it is much faster to use SAMtools (Li et al., 2009) or another tool to extract the homoeologous region of interest before running CAPG.

##### S1.1.1 Data preprocessing

**Selection of homoeologous regions.** CAPG is intended for genotyping homoeologous regions; regular diploid genotypers can be used for non-homoeologous regions and updog (Gerard et al., 2018) can be used for regions with incomplete preferential pairing or completely autopolyploidized regions. To identify homoeologous regions for genotyping, one may use a local aligner, such as BLAST, or a global aligner, such as MUMmer4 (Marçais et al., 2018), to align the subgenomic reference genomes. We used BLAST for our small-scale validation study. The selected regions are communicated to CAPG as `CHROM:START-END`, where `CHROM` is the name of a chromosome in the subgenomic reference and `START` and `END` are the 1-based inclusive termini of the targeted region in that chromosome.

**Genome alignment files.** Once the homoeologous regions have been selected, CAPG requires a SAM-formatted alignment of the subgenome reference sequences in each homoeologous region. Such a file can be generated with MUMmer4 or for sufficiently short regions, any software for global pairwise alignment. We used MUMmer4 with default parameters. CAPG uses the QNAME, FLAG, RNAME, POS, and CIGAR fields, but not the SEQ or QUAL fields, so it is not affected by software that does not reverse complement the SEQ or reverse the QUAL when the FLAG indicates it.

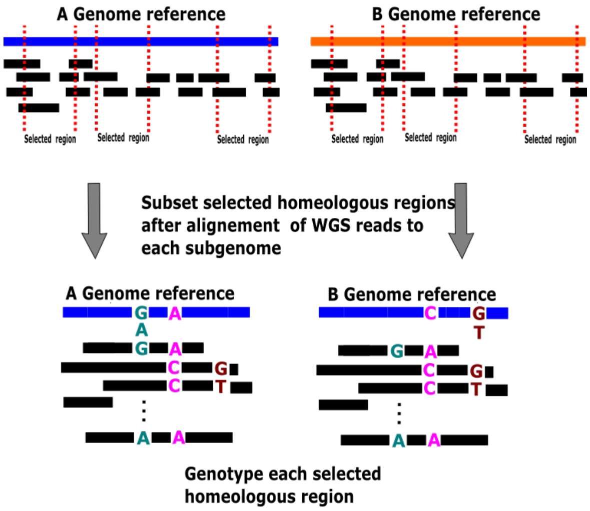

Figure S1: Pre-processing WGS resequencing data. Reads aligning to any portion of a selected homoeologous region are extracted and used for genotyping the loci within that region.

**Read alignment.** Reads are aligned to each subgenomic reference. Any short read reference alignment tool, excluding pseudoaligners, can be used to perform whole genome alignments of the sequence reads to each of the subgenomic references. In this study we have used BWA-MEM (Li et al., 2009) with default parameters.

**Extraction of relevant reads.** Given the whole genome read alignments and the bounds of the selected homoeologous regions, Figure S1 shows how all reads overlapping the selected regions are collected for genotyping. CAPG takes two SAM files for these alignments and uses QNAME, FLAG, RNAME, POS, CIGAR, SEQ and QUAL fields. It assumes SEQ is reverse complemented and QUAL reversed when the FLAG indicates it. CAPG also needs FASTA files containing the subgenomic references in order to evaluate the alignment of reads that extend beyond the target regions.

**Read quality.** After alignment, the user may wish to discard reads with likely errors. By default, CAPG only discards unmapped reads, supplementary alignments, and reads that do not align exactly once to each homoeolog, but the user may also wish to discard reads with excessive soft-clipping or indels, as well as reads that are too short, too long, or of low quality. Such reads may contain more noise than useful information. It is also possible to discard secondary alignments, genotyping from the surviving primary alignments, but since such reads may be misplaced from repetitive or paralogous loci, retaining only the primary alignments may increase false evidence for heterozygosity.

**Minimum coverage.** In the selected homoeologous region, we use all $n_k$ reads that have homoeologous alignments in both subgenomes to the locus of interest in individual $k$ for genotyping. CAPG will genotype the locus as long as $n_k > 0$, but a user may wish to impose a coverage requirement. Using the read likelihoods of the selected reads, CAPG can impose a minimum requirement on the expected coverage

$$E_{ks} = \sum_{i=1}^{n_k} \Pr(S_i = s \mid \boldsymbol{r}_i) \tag{S1}$$

of subgenome $s$ at the locus, assuming homozygous reference genotypes at every read position with homoeologous alignment except the current locus, which is ignored, and equal subgenomic coverage $\Pr(S_i = s) = 0.5$ for $s \in \{1, 2\}$. Alternatively, the user may proceed to genotype with CAPG and impose a coverage filter later in the analysis pipeline.

**Other screens.** Since our tool is not intended to genotype indel variants, we skip sites with homoeologous indels in the subgenomic reference alignment. Meanwhile, segregating indels can be detected and screened when the expected subgenomic coverage $E_{kls}$, defined in Eq. (S1), falls substantially below the expected subgenomic coverage of all reads covering the locus, including those without homoeologous alignments. Users may also wish to skip genotyping a locus when there is a third allele with coverage exceeding a user-provided percentage (default: 50%) of the smaller expected subgenomic coverage. Alleles reaching this level of coverage are likely true variants, but CAPG assumes biallelic SNPs.

##### S1.1.2 Post-genotyping filters

**Equal homologous coverage test.** Our genotyping likelihood assumes all $n$ reads (dropping index $k$ for individual to simplify notation) used to genotype the locus are sampled with equal read coverage of all four chromosomes, but we can fit a model that allows unequal coverage. Specifically, we modify the $i$th read likelihood, previous Eq. (2), to

$$\begin{aligned}\Pr(\boldsymbol{R}_i = \boldsymbol{r}_i \mid \boldsymbol{M}) = \eta \Pr(R_{ij_i} = r_{ij_i} \mid S_i = 1, M_1 = m_1) \prod_{\substack{j=1 \\ j \neq j_i}}^{l_i} \Pr(R_{ij} = r_{ij} \mid S_i = 1) \\ + (1-\eta) \Pr(R_{ij_i} = r_{ij_i} \mid S_i = 2, M_2 = m_2) \prod_{\substack{j=1 \\ j \neq j_i}}^{l_i} \Pr(R_{ij} = r_{ij} \mid S_i = 2),\end{aligned}$$

where $\eta$ is the probability of sampling a read from subgenome A, and the probability of observing nucleotide $r_{ij_i}$ aligned to the locus is

$$\Pr(R_{ij_i} = r_{ij_i} \mid S_i = s, M_s = 1) = \Pr(R_{ij_i} = r_{ij_i} \mid T_i = t_r)\,\gamma_s + \Pr(R_{ij_i} = r_{ij_i} \mid T_i = t_a)\,(1-\gamma_s),$$

where $\gamma_s$ is the probability of sampling a reference allele from subgenome $s$ when subgenome $s$ is heterozygous. When subgenome $s$ is homozygous, then $\gamma_s \in \{0, 1\}$ is not a free parameter. Furthermore, for heterozygous subgenome $s$ ($M_s = 1$), the null hypothesis of equal homolog coverage is $H_0 : \gamma_s = 0.5$. We perform a likelihood ratio test of this nested $H_0$ against the alternative hypothesis $H_a : \gamma_s \in [0, 1]$. We use an EM algorithm to maximize the likelihood over $\eta$ to allow unequal subgenomic coverage. The resulting $p$-value can be used to identify suspicious heterozygous calls.

**Equal homoeologous coverage test.** One can also test the hypothesis of equal homoeologous coverage, $H_0 : \eta = 0.5$, to indicate unusually biased subgenomic coverage. The model described above is fit with $\gamma_s$ determined by the genotype estimate $\hat{\boldsymbol{M}}_k$ or estimated if $\hat{M}_{ks} = 1$. In particular, one explanation for a high quality PCR error, say A, in otherwise monomorphic reads, say T, when there are few nearby homoeologous SNPs is to hypothesize the error exists as genotype AT in one subgenome observed with low coverage. While this explanation does not lead to a low $p$-value in the equal homologous coverage test, it can still be detected as unusually biased homoeologous coverage. However, since homoeologous coverage may legitimately vary more than homologous coverage, one may wish to apply a more stringent threshold on the $p$-value or threshold on $\hat{\eta}_s$ instead. Furthermore, since both these tests require running an iterative EM algorithm, they will slow down genotyping, especially for the equal homoeologous coverage test, which can be applied to every locus, rather than just the loci with heterozygous calls.

**Identifiability.** Mismatches between the subgenome references that are true homoeologous SNPs are key to correctly assigning reads to their source subgenome. If there are no mismatches within read distance of the locus to genotype (nonidentifiable loci), then CAPG has little information to source any reads covering the locus and will produce ambiguous posterior probabilities (1). (A little information can come from nearby heterozygous loci, where the coverage of the alternative allele should be a fourth of the total coverage under the assumption of uniform coverage of all four chromosomes.) GATK can successfully detect such loci as homoeologous SNPs since the short read aligner may be able to use the subgenome mismatch to partition reads. We have discussed this phenomenon in the main text and will discuss it further in §S2.2.1. In some contexts, it makes sense to discard nonidentifiable loci. Though it is possible, and more effective, to discard nonidentifiable loci as a pre-genotyping filter that checks for reads covering both the current locus and nearby subgenomic mismatches, we have filtered them in this study in post-processing by checking for subgenomic mismatches within read length distance of the current locus.

| Simulation | Coverage $c$ | Homoeologous Rate $r_h$ | No. of Loci with No Coverage In Individuals | Across Sample |
|---|---|---|---|---|
| 1 | 10 | 0.007 | 1413 | 2 |
| 2 | 20 | 0.007 | 849 | 0 |
| 3 | 40 | 0.007 | 386 | 0 |
| 4 | 40 | 0.005 | 369 | 0 |
| 5 | 40 | 0.010 | 362 | 0 |

Table S1: **Properties of simulated datasets.** The coverage rate $c$ is the expected number of reads covering each locus in each subgenome. The homoeologous rate $r_h$ is the probability of a homoeologous SNP at each locus. We report the number of loci with no covering reads in individuals and the number of loci with no coverage in *any* individual in the sample.

#### S1.2 GATK metrics

For genotyping a locus, our procedure has split the reads into disjoint subsets of $n_{kg}$ reads $\mathcal{R}_{kg} = \{\boldsymbol{r}_{kg1}, \boldsymbol{r}_{kg2}, \ldots, \boldsymbol{r}_{kgn_{kg}}\}$ with a base aligned to the locus of subgenome $g \in \{0, 1\}$ in the $k$th individual. The normalized Phred-scale likelihood (PL) is a sample-level (per individual) annotation calculated by HaplotypeCaller, recorded in the sample-level columns of variant records in VCF files. It is defined as

$$\mathrm{PL}_{klg}(m) = -10\big[\log_{10} \Pr(M_{kg} = m \mid \mathcal{R}_{kg}) - \log_{10} \Pr(M_{kg} = \widehat{M}_{kg} \mid \mathcal{R}_{kg})\big] \tag{S2}$$

for the genotypes considered in the variant record for each sample. Furthermore, the allotetraploid genotype $\boldsymbol{m} = (m_1, m_2)$ has normalized likelihood

$$\mathrm{PL}_k(\boldsymbol{m}) = \mathrm{PL}_{k1}(m_1) + \mathrm{PL}_{k2}(m_2),$$

by independence of the genotype calls within each subgenome. Because the second term in Eq. (S2) is a constant, it cancels when computing differences of PL values at any locus in subgenome $g$ of individual $k$.

The metrics supporting heterozygosity, allelic SNPs, homoeologous SNPs, Eqs. (4)–(6), in terms of PL values, become

$$0.1 \ln(10) \left[ \min_{m \in \{0,2\}} \mathrm{PL}_{kg}(m) - \mathrm{PL}_{kg}(1) \right] \tag{S3}$$

$$0.1 \ln(10) \min_{\mathrm{N} \in \{\mathrm{N}_1, \mathrm{N}_2\}} \sum_{k=1}^{n} \big[ \mathbb{1}_{\{t_{kr} = \mathrm{N}\}} \mathrm{PL}_{kg}(0) + \mathbb{1}_{\{t_{ka} = \mathrm{N}\}} \mathrm{PL}_{kg}(2) \big] \tag{S4}$$

$$-0.1 \ln(10) \min_{\mathrm{N} \in \{\mathrm{N}_1, \mathrm{N}_2\}} \sum_{k=1}^{n} \big[ \mathbb{1}_{\{t_{kr} = \mathrm{N}\}} \mathrm{PL}_{k}(0,2) + \mathbb{1}_{\{t_{ka} = \mathrm{N}\}} \mathrm{PL}_{k}(2,0) \big]. \tag{S5}$$

As for CAPG, the PL value of the required genotype may not be available for some loci and individuals, in which case we may substitute the maximum PL value reported for the individual at the locus.

#### S1.3 Simulation procedure

A 10,000bp-length region of subgenome A chromosome Arahy.chr01 from the Tifrunner peanut assembly (Bertioli et al., 2016) was selected as a reference for simulation. To simulate a "homoeologous subgenome B", we independently mutate each site of the selected region with probability $r_h$, at an average 10,000×$r_h$ homoeologous sites. The substituted nucleotide is any of the remaining three with equal probability. To simulate allelic SNPs, we assume HWE in the population, and then sample allelic proportions from a Beta($\alpha, \beta$) distribution with $\alpha = \beta = 100$. We independently select non-homoeologous sites with probability $2r_a$, choose the affected subgenome with equal probability, and then select an alternate allele with equal probability. After these modifications, each site is either a homoeologous SNP, a subgenome A allelic SNP, a subgenome B allelic SNP, or conserved (monomorphic); and there are no more than two nucleotides per site. For the $k$th simulated individual, we independently (assuming linkage equilibrium) sample a genotype for each allelic SNP and finalize the four haplotypes. Lastly, we generate reads for the $k$th individual using ART (Huang et al., 2012). A summary of the simulated datasets is shown in Table S1. Error profiles were estimated using the ART

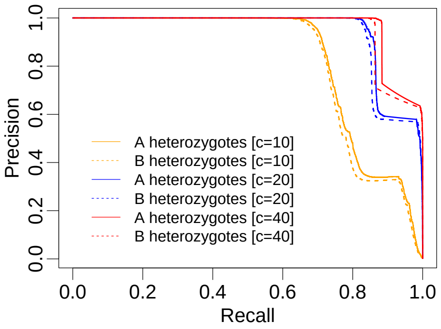

(a) Heterozygosity, coverage varying.

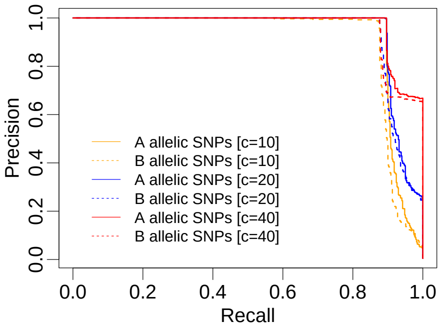

(b) Allelic SNPs, coverage varying.

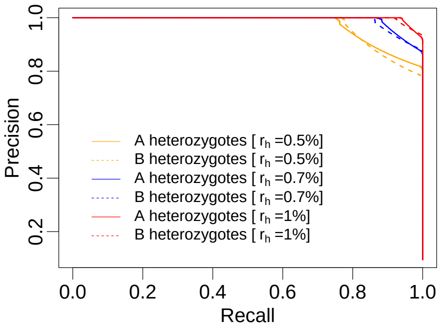

(c) Heterozygosity, $r_h$ varying.

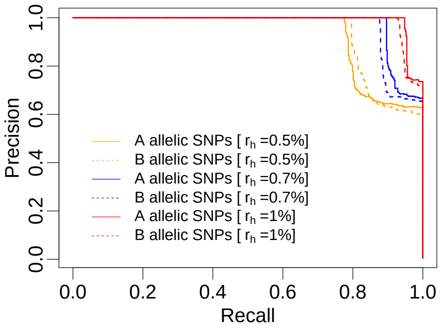

(d) Allelic SNPs, $r_h$ varying.

Figure S2: **PR curves for CAPG on simulated allotetraploid data.** Expected coverage $c$ per subgenome varies with fixed homoeologous rate per site $r_h = 0.007$ in (a) and (b). Homoeologous rate $r_h$ varies with fixed coverage $c = 40$ in (c) and (d). Support for allelic SNPs evaluated with Eq. (5) in (b) and (d); support for heterozygosity evaluated with Eq. (4) in (a) and (c).

Illumina profiler applied to 150bp MiSeq paired end reads from WGS resequencing data of peanut genotype OLin (Clevenger et al., 2017).

Thanks to the suggestion of a reviewer, we also implemented a second simulation to mimic the scenario where one of the subgenomic reference sequences is of lower quality than the other. For this simulation, we randomly introduced nucleotide mismatches, assuming uniform probability over the possible nucleotides, at proportion $r_m \in \{0, 0.01, 0.1\}$ of the sites in the subgenome A reference used for alignment (but not simulation), while holding $r_h = 0.007$ and $c = 10$. We simulated the $r_h = 0.007$, $c = 10$ and $r_m = 0$ read data again because we had not retained it from the original simulation, which explains differences with previous PR curves at this simulation setting.

### S2 Results

#### S2.1 Simulation

We evaluated how well CAPG predicts heterozygosity, allelic SNPs, and homoeologous SNPs on simulated allotetraploid data with varying coverage ($c$) and homoeologous rate ($r_h$). Precision-Recall (PR) curves in Figures S2a and S2c show improved prediction of heterozygotic loci with increasing coverage or homoeologous rate. The area under the PR curve (AUCPR) increases from 0.82 to 0.96 as coverage climbs from 10 to 40 reads per subgenome, with $r_h = 0.007$ fixed (Table S2). AUCPR climbs from 0.89 to 0.99 as homoeologous rate increases from 0.5% to 1%, with coverage $c = 40$ fixed. PR curves in Figures S2b and S2d also show improved prediction of allelic SNPs with increasing coverage (Table S2, rows 1–3: AUCPR increases from 0.91 to 0.97) or homoeologous rate (Table S2, rows 4–6: AUCPR increases from 0.93 to 0.99). Increasing coverage also improves homoeologous SNP detection (Table S2, rows 1–3) as does increasing homoeologous rate (Table S2, rows 4–6), but with baseline levels already so high, the improvement is small and PR curves are not shown.

As explained in the main text, the superior performance of GATK for calling allelic SNPs is a consequence of overcalling heterozygosity on nonidentifiable loci without nearby homoeologous SNPs. After removing such nonidentifiable loci, Table S2 (rows 7–12) show CAPG has better performance than GATK on the remaining loci.

| | | **CAPG** | | | | | **GATK** | | | | |
|---|---|---|---|---|---|---|---|---|---|---|---|
| | | **Heterozyg.** | | **Allelic** | | **Homeo-** | **Heterozyg.** | | **Allelic** | | **Homoe-** |
| $r_h$ | **Cov.** | **A** | **B** | **A** | **B** | **logous** | **A** | **B** | **A** | **B** | **logous** |
| | | | | | | **All Loci** | | | | | |
| 0.7% | 10 | 0.83 | 0.82 | 0.92 | 0.91 | 0.99 | 0.49 | 0.52 | **0.97** | **0.98** | 0.98 |
| 0.7% | 20 | 0.94 | 0.93 | 0.94 | 0.93 | 1.00 | 0.66 | 0.67 | **0.98** | **0.99** | 0.98 |
| 0.7% | 40 | 0.96 | 0.95 | 0.97 | 0.96 | 1.00 | 0.71 | 0.71 | **0.98** | **0.99** | 0.98 |
| 0.5% | 40 | 0.89 | 0.90 | 0.93 | 0.93 | 1.00 | 0.59 | 0.58 | **0.95** | **0.95** | 0.79 |
| 0.7% | 40 | 0.96 | 0.95 | 0.97 | 0.96 | 1.00 | 0.71 | 0.71 | **0.98** | **0.99** | 0.98 |
| 1.0% | 40 | 0.99 | 0.98 | 0.99 | 0.98 | 1.00 | 0.83 | 0.82 | **0.99** | **1.00** | 0.99 |
| | | | | | | **Unidentifiable Loci Removed** | | | | | |
| 0.7% | 10 | 0.88 | 0.89 | 1.00 | 1.00 | 0.99 | 0.57 | 0.60 | 0.99 | 0.99 | 0.98 |
| 0.7% | 20 | 0.98 | 0.99 | 1.00 | 1.00 | 1.00 | 0.75 | 0.76 | 0.99 | 1.00 | 0.98 |
| 0.7% | 40 | 1.00 | 1.00 | 1.00 | 1.00 | 1.00 | 0.82 | 0.82 | 0.99 | 1.00 | 0.98 |
| 0.5% | 40 | 0.99 | 0.99 | 1.00 | 1.00 | 1.00 | 0.77 | 0.76 | 0.99 | 0.99 | 0.97 |
| 0.7% | 40 | 1.00 | 1.00 | 1.00 | 1.00 | 1.00 | 0.82 | 0.82 | 0.99 | 1.00 | 0.98 |
| 1.0% | 40 | 1.00 | 1.00 | 1.00 | 1.00 | 1.00 | 0.89 | 0.89 | 0.99 | 1.00 | 0.99 |

Table S2: AUC values for PR curves for simulation data without removing loci (**top**) and after removing nonidentifiable loci with no homoeologous SNPs within read distance (**bottom**). AUCPR values are highlighted wherever GATK is equal or superior to CAPG. **Cov.** is expected subgenomic coverage. **Heterozyg.** is for genotyping heterozygotes.

In a final simulation, we varied the rate of mismatches $r_m \in \{0, 0.001, 0.01\}$ in the subgenome A reference used for alignments to explore the cost of using a lower quality reference sequence for one of the subgenomes. The homoeologous SNP rate $r_h = 0.007$ is above all but the last mismatch rate,

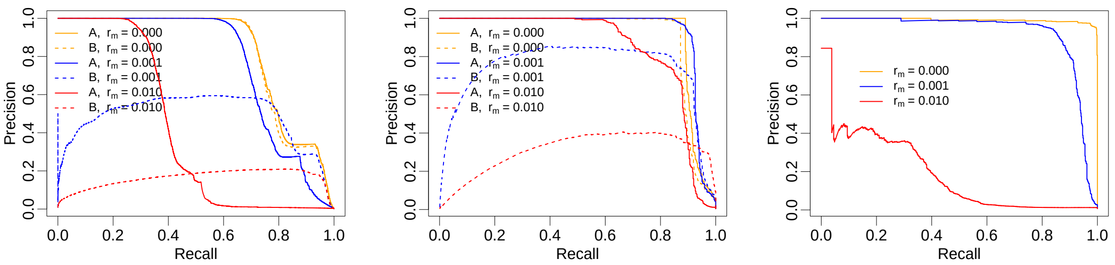

(a) Heterozygous genotypes (b) Allelic SNPs (c) Homoeologous SNPs

Figure S3: Poor reference quality hurts genotyping performance. We introduce additional mismatches in the reference for subgenome A to mimic the use of a reference genome distantly related to the actual subgenome. We uniformly substitute to any of the three other nucleotides at rates $r_m = 0.01$ or $r_m = 0.10$ and compare PR curves with results obtained when using a reference with no additional variation.

and coverage $c = 10$ is low. The excess mismatches in the subgenome A reference will not match the reads except by error. Fig. S3 shows that increasing mismatch rate $r_m$ progressively decreases the performance, preferentially disrupting the calls in the high quality genome. The substituted loci in the subgenome A reference drive reads to increasingly be assessed as more likely products of the subgenome B reference, regardless of their true source. Average coverage of subgenome A drops from 9.8, equal to subgenome B coverage when there are no mismatches, to 8.9 with $r_h = 0.001$ and 4.6 with $r_h = 0.010$. The rogue subgenome A reads assigned to subgenome B contaminate genotyping in the latter subgenome. It might be possible to detect the problem by requiring loci to meet a minimum expected coverage of both subgenomes via Eq. (S1) or pass the equal homoeologous coverage test of §S1.1.2. It would also be possible to correct the subgenome A reference when the reads indicate a monomorphic site.

#### S2.2 Real data

Tables S3 and S4 show the peanut and cotton genotypes and accessions examined in our analysis of real data.

For final genotype and SNP calling, loci were filtered for an expected subgenomic coverage of at least eight in both subgenomes and heterozygous calls were discarded if the equal homologous coverage hypothesis was rejected at significance level 0.05. In peanut, 1,4008,313 of 44,268,979 loci (31%) were removed for low coverage. Of 47,807 heterozygous calls at high coverage loci, 36,962 (90%) were discarded for failing the equal homologous coverage test. These filters prevented SNP calling at 54,251 of 3,169,541 loci (1.7%) by eliminating data from all genotypes. Since we always applied these filters to the cotton data, we do not have these numbers for cotton. The histograms of the three CAPG metrics (4)–(6) for the surviving loci in peanut are shown in Figure S4(d)–(c) along with the thresholds chosen for calling heterozygosity ($> 80$), as well as allelic ($> 5$) and homoeologous SNPs ($= 0$). For comparison, histograms of the metrics from the two simulation conditions with closest subgenomic coverage, 10 in S4(g)–(i) and 20 in S4(j)–(l), are also shown. After applying the chosen thresholds, the counts of each type of call are reported in the main text Tables 1 and 2.

| Genotype | SRA accession | Genotype | SRA accession |
|---|---|---|---|
| Florida07 | SRR4124062 | HRP32 | SRR8361736 |
| KatieSARI | SRR4124066 | HRP31 | SRR8361737 |
| COC230 | SRR4124068 | HRP34 | SRR8361738 |
| Florunner | SRR4124074 | ICG2857 | SRR8736998 |
| NMValencia | SRR4124078 | ICG4527 | SRR8737008 |
| HRP36 | SRR8361734 | ICG12370 | SRR8737061 |
| HRP35 | SRR8361735 | ICG9362 | SRR8737062 |

Table S3: List of peanut accessions selected for the real data analysis.

| Genotype | SRA accession | Genotype | SRA accession |
|---|---|---|---|
| Coker139 | SRR1580587 | Yucatanense 6 | SRR1580671 |
| Stoneville2B | SRR1580604 | *Richmondi 3* | SRR1580675 |
| Xinluzao 26 | SRR1580630 | Marie-galante 71 | SRR1580682 |
| Punctatum_25 PE | SRR7887417 | CIR_12 PE | SRR7887414 |
| Brazil cotton | SRR1580642 | | |

Table S4: List of cotton accessions selected for the real data analysis.

##### S2.2.1 Comparing CAPG and GATK without a known truth

With no truth to compare against, we examined relationships between the metrics and other data characteristics at genotyped loci in the peanut data (Table S5 and Figures S7–S9) to try and assess method accuracy. For this analysis, we have applied no filters. The CAPG and GATK metrics for heterozygosity are well-correlated (Spearman correlation > 0.9, row CAPG in Table S5), but much less correlated for SNP calls (Spearman correlation < 0.4), thus confirming a substantial difference between the methods. We took three approaches to compare CAPG and GATK metrics: (1) inspection of alignments and manual genotyping, (3) an assessment whether nonidentifiable loci, physically impossible to genotype, affect the computed metrics, and (2) a comparison of the genotype calls to a rough truth inferred from the existing mismatches between the subgenomic references. We examine each of these lines of evidence next.

First, we inspected 22–64 loci with varying values for each metric to manually genotype the individuals and assign a "truth". Read aligments for an example showing a C/G homoeologous SNP are shown in Figure S5. There is a subgenomic mismatch 33bp 5' and 17bp 3' of the SNP locus, but the 5' mismatch is monomorphic in all accessions we genotyped. Since CAPG does not use the information from the genotyped locus to assign reads to a subgenome, the false information from the 5' locus balances nearly perfectly the information from the 3' homoeologous SNP for reads traversing both sites. The 5' mismatch drives four of 18 subgenome B reads to subgenome A in SRR8361736 and nearly 9 of 21 reads in SRR8361737. The correct genotype CC/GG is called in SRR8361736, but CG/GG is called in SRR8361737. The allelic SNP metric is 80.4 for CAPG and 0 for GATK. The upstream mismatch is correctly called monomorphic by both methods, suggesting correction of the subgenomic references followed by a second round of genotyping could eliminate the CAPG miscall.

Overall, CAPG metrics are more associated with hand-verified calls of allelic SNPs (Pearson correlation 0.771 for CAPG vs. 0.685 for GATK) and homoeologous SNPs (Pearson correlation 0.828 for CAPG and 0.695 for GATK). There was no assessment for the heterozygous metric because we could not unambiguously confirm any heterozygous call. While the sampled loci were not selected at random, we sampled loci from all four quadrants of the scatter plots (Figure 5), with the following caveats. We repeatedly sampled from the lower right of the scatter plot of allelic SNP metrics, where the CAPG metric is high and the GATK metric is low, but we could not discern the truth for most of these loci. These loci tend to be in regions with many mismatches between the subgenomes, some of them monomorphic in the sampled genotypes, and additional sites of variation where the subgenomic references match. Frequently there was evidence of more than four haplotypes, suggesting reads aligning from paralagous sources. It seems reasonable to expect strong allelic SNP signal in such regions, but there is no truth to confirm and these loci are left unresolved ("Unclear" in Figures 5 and S6). Of the four loci from this quadrant where a truth could be determined, the site was assessed to be a homoeologous SNP, at odds with the CAPG metric. In two cases, where the transformed CAPG metric was below 0, the locus was distant from homoeologous SNPs and the CAPG metric reflects uncertainty about the true call, whether an allelic or homoeologous SNP. As demonstrated in Figure S5, GATK confidently calls homoeologous SNP because the aligner has used the locus in question to assign reads to subgenomes. We would argue that the CAPG metric better reflects a valid uncertainty in these genotype calls. The other two loci, one of which was examined in Figure S5, have a nearby subgenomic mismatch that is monomorphic in the sampled genotypes. As suggested above, such false calls should resolve after correcting the subgenomic references. We also examined but did not type loci falling in the lower right of the homoeologous scatter plot. All of these loci have no more than one expected read in at least one subgenome and are completely and sensibly removed by the coverage filter in practice. In the upper left of both plots, where GATK favors a SNP call and CAPG does not, we found evidence of monomorphic sites miscalled by GATK.

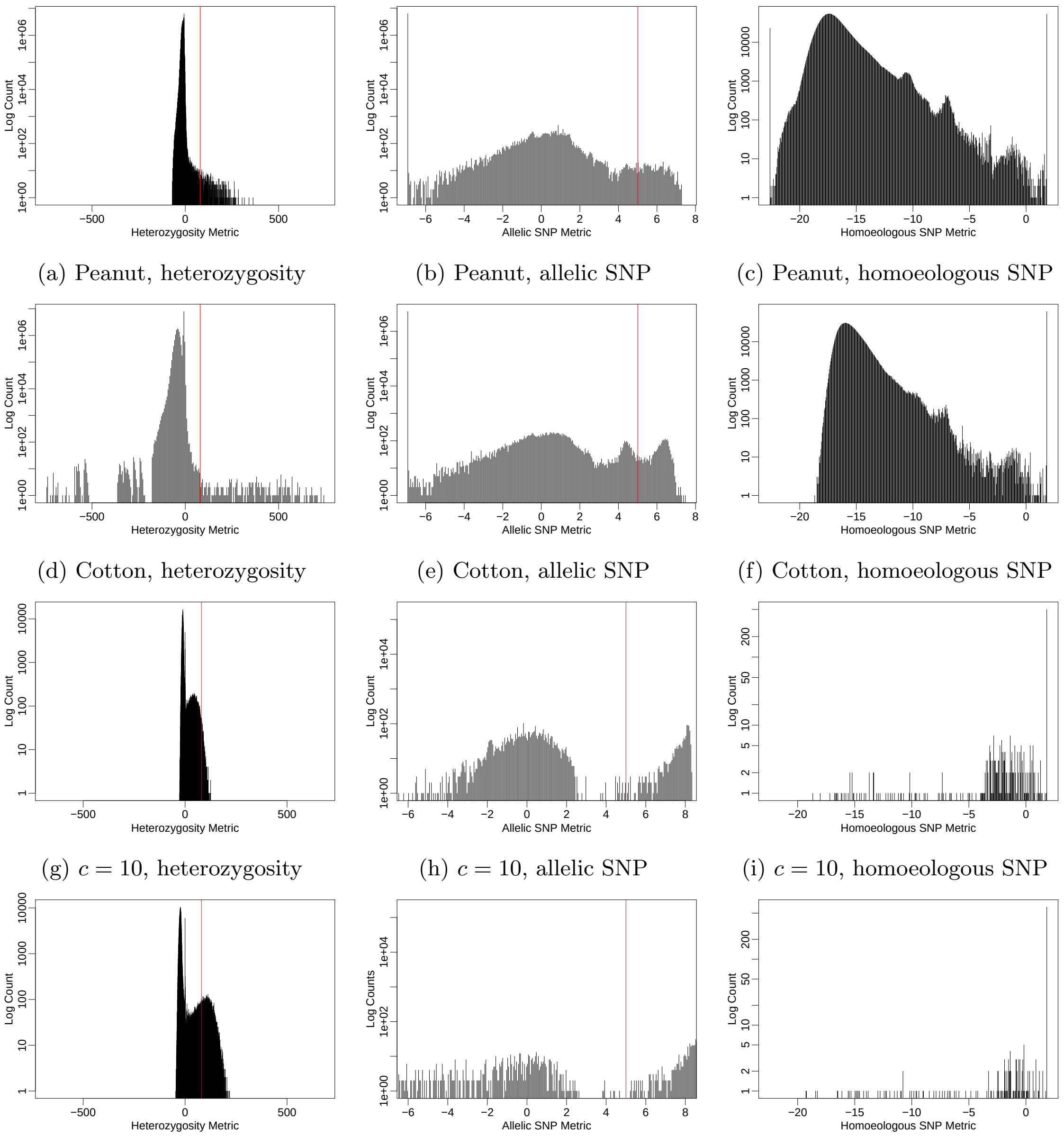

(a) Peanut, heterozygosity (b) Peanut, allelic SNP (c) Peanut, homoeologous SNP

(d) Cotton, heterozygosity (e) Cotton, allelic SNP (f) Cotton, homoeologous SNP

(g) $c = 10$, heterozygosity (h) $c = 10$, allelic SNP (i) $c = 10$, homoeologous SNP

(j) $c = 20$, heterozygosity (k) $c = 20$, allelic SNP (l) $c = 20$, homoeologous SNP

Figure S4: Histograms of heterozygosity metric (4) (**first column**), transformed allelic SNP metric (5) (**second column**) and transformed homoeologous SNP metric (6) (**third column**) from peanut data (**first row**), cotton data (**second row**) simulation data with coverage $c = 10$ (**third row**), and simulation data with coverage $c = 20$ (**fourth row**). For real data, these are the metrics after locus filtering as described in the text. Heterozygosity metrics and allelic SNP metrics from both subgenomes are combined in these plots. Allelic SNP metrics are log transformed after adding $1 \times 10^{-3}$; homoeologous SNP metrics are subject to the Box-Cox transformation discussed in legend of Fig. 3. The $y$-axis is log transformed to overcome massive excess of monomorphic loci with low values of the metrics. The thresholds we utilized to call heterozygosity (80) and allelic SNPs (5) in the real data are shown in red. Homoeologous SNPs were called when the metric was equal to 0, where a point mass is clearly visible in the plots.

We next determined whether genotype calling is possible at a locus, *i.e.*, whether reads at the locus can be sorted into subgenomes. Specifically, we labeled all loci with at least one mismatch between the subgenomic references within read distance (150bp) of a locus as "identifiable." A genotyper should produce ambiguous metrics at nonidentifiable loci, and generally more extreme, decisive metrics at identifiable loci. We assess increased spread in the distribution of metrics at identifiable loci using Levene's test (Levene, 1960). Both genotypers produce metrics at least somewhat sensitive to identifiability, but for heterozygous and allelic SNPs CAPG is more sensitive than GATK (Identifiable, Table S5). GATK is slightly more sensitive to identifiability when calling homoeologous SNPs, but it identifies homoeologous SNPs when they are not identifiable by our definition (upper, right subplot of Figure S9b) because the genotyped site is used to partition the reads between subgenomes.

Lastly, mismatches between the subgenomic references could be homoeologous SNPs, allelic SNPs or errors. As such, they represent a fuzzy indication of the truth of homoeologous SNPs and allelic SNPs, and a good genotyper will tend to produce higher SNP metrics for mismatches. The hypothesis is confirmed for both genotypers in Figures S8 and S9, where the metric distributions for mismatching loci (purple) shift right relative to the distribution for matching loci (green). The association is stronger for CAPG on allelic SNPs in both genomes, but slightly stronger for GATK on homoeologous SNPs (Mismatch, Table S5), another artefact of using the genotyped site to assign reads to subgenomes. The distortion in the GATK homoeologous metric is misleading. While CAPG can find some non-mismatch sites it calls homoeologous SNPs, GATK does not (Figure S9). Further, GATK finds relatively fewer mismatch sites to be allelic SNPs, even though some of the reference mismatches must be captured allelic differences (Figure S8).

Using the presence of mismatch as a noisy "truth" label for allelic or homoeologous SNPs further demonstrates these observations (PR curves in Figure S10). GATK predicts all mismatches to be homoeologous SNPs, but CAPG finds strong homoeologous SNP support for less than 90% of mismatches. Another fraction of the subgenomic mismatches are likely allelic SNPs, and far more of these find support from CAPG than GATK (Figure S10b), but the majority of allelic SNPs are found at matching sites by both CAPG and GATK. It would be concerning if many allelic SNPs were captured as subgenomic mismatches since such sites disrupt read alignments and alignment likelihoods.

There are some additional strong correlations between the metrics and data characteristics that are to be expected. Subgenomic coverage is strongly and negatively correlated with the heterozygosity metrics (Coverage, Table S5 and Figure S7 for subgenome A), because higher coverage increases confidence in the call, which is non-heterozygous, thus a low metric, for the vast majority of loci. The sum of average subgenome coverage also negatively correlates with the homoeologous metrics, though much more strongly for GATK (Spearman correlation $-0.796$) than CAPG ($-0.485$). The average subgenome coverage is much less strongly correlated with the allelic metrics ($-0.05$ to $-0.07$), probably because it is the coverage of particular individuals, such as heterozygotes, not all individuals, that determines the strength of the evidence for allelic SNPs. It is likely the previously mentioned correlations, plus a strong correlation in coverage of the homoeologous loci across subgenomes (not shown), that induces a positive correlation in the heterozygosity and allelic metrics across the two subgenomes (Metric B, Table S5).

##### S2.2.2 Posthoc filtering

Filtering by low coverage or the equal homologous coverage test (EHCT) removes genotype calls at some loci, overall reducing evidence for allelic SNPs and increasing evidence for homoeologous SNPs (Figure S6) and monomorphic loci. Genotyping fewer individuals obviously decreases the chance to discover allelic SNPs (Nielsen et al., 2011), but the EHCT particularly targets and eliminates heterozygous calls that would otherwise support allelic SNPs. Filtering removes a portion of the false heterozygous genotypes in GATK and weakly supported heterozygous genotypes in both methods (compare unfiltered metrics in Figure 4 to filtered metrics in Figure S6a). Filtered loci are more likely to be called homoeologous SNPs, including three manually confirmed homoeologous SNPs, and less likely to be called allelic SNP, including one hand-verified allelic SNP, in both CAPG and GATK. The most severe shifts in support for allelic SNPs, however, affect already borderline calls (bottom row of points in Figure S6). Overall, the coverage and EHCT filtering neither eliminates the disagreements between methods nor universally fixes the incorrect decisions assessed by manual genotyping.

Additional filters may further improve genotyping and SNP calling. We proposed but did not apply a test of equal homoeologous coverage (§S1.1.2), which may help to filter false heterozygous

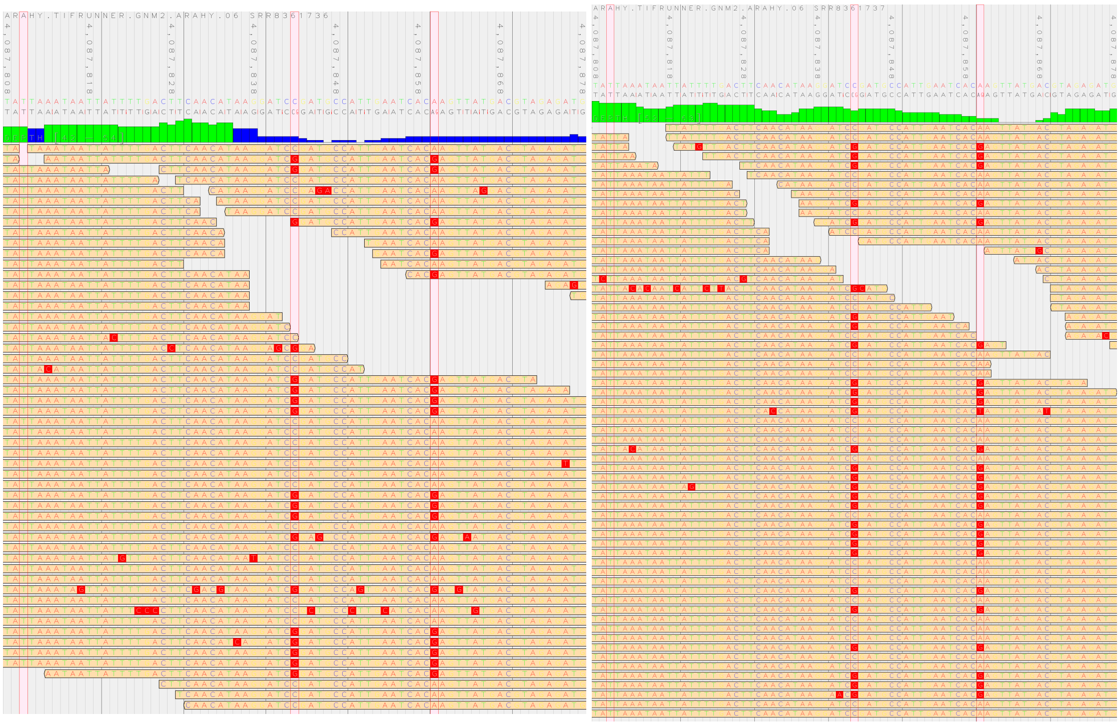

(a) SRR8361736

(b) SRR8361737

Figure S5: Pileup of reads covering site 4,087,843 (center highlighted locus, outlined in red) of chromosome 6 (homoeologous locus 18,944,270 in chromosome 16) in peanut accessions (a) SRR8361736 and (b) SRR8361737 visualized using JVarKit (Lindenbaum, 2015). Reads are shown aligned to subgenome A, so the alternate alleles shown on a red background are errors or alleles present in subgenome B. There is a subgenomic mismatch 33bp (left highlighted locus) before the site and 17bp (right highlighted locus) after the site, but only the latter is variable in these two accessions (and all accessions). Although the two SNPs are in complete linkage disequilibrium, CAPG cannot place the reads while ignoring the information from the center locus to be genotyped. It calls SRR8361736 CC/GG and SRR8361737 CG/GG. Meanwhile, GATK uses the information from the center locus and correctly genotypes the locus as CC/GG.

calls triggered by a small number of high quality errored reads, perhaps due to PCR amplification or paralogous read mapping. Paralogous reads can manifest as (1) more than four total haplotypes covering a site or (2) atypical coverage levels for an allele not in the set $\{0, 0.25, 0.50, 0.75, 1\}$. The equal homologous and homoeologous coverage tests can detect atypical coverage levels, but both manifestations were used to screen reads in a version of CAPG for amplicon data, where off-target amplification of paralogs is a serious problem. See https://github.com/Kkulkarni1/CAPG.git for more information.

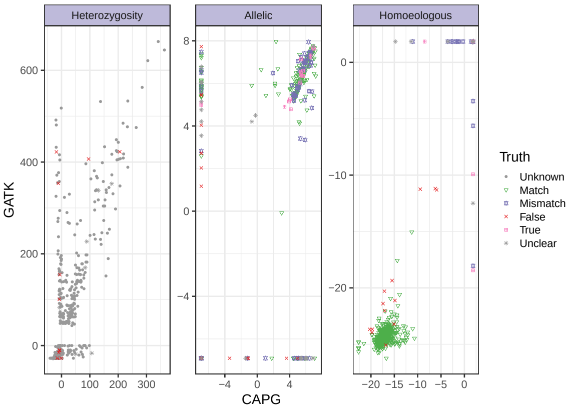

(a) All, filtered

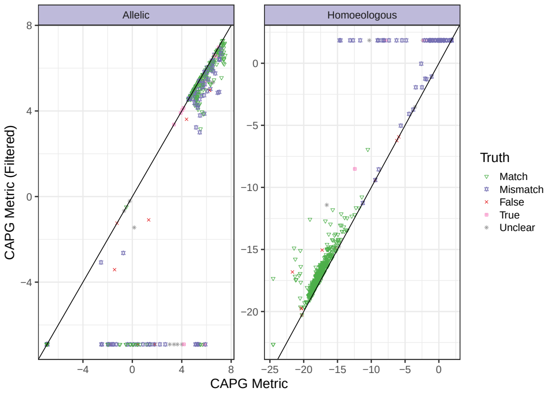

(b) CAPG SNP calls

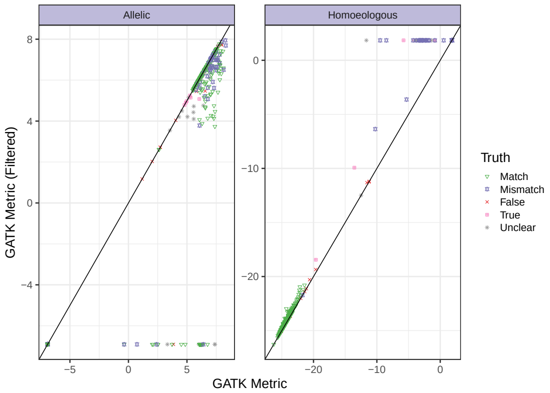

(c) GATK SNP calls

Figure S6: Affect of coverage and equal homologous coverage test (EHCT) filtering on the SNP metrics. Scatter plot of CAPG vs. GATK metrics after filtering (a). Compare unfiltered heterozygosity metrics in Figure 4 and see the legend there for more information. For SNP calls, we also plot allelic SNP metrics (left facet) and homoeologous SNP metrics (right facet) for CAPG (b) and GATK (c) before ($x$-axis) and after ($y$-axis) applying the coverage and ECHT filters.

| | CAPG | | GATK | |
|---|---|---|---|---|
| | **Statistic** | **$p$-value** | **Statistic** | **$p$-value** |
| **Heterozygous A** | | | | |
| Identifiable | 50 643 | 0 | 145 | $2.6 \times 10^{-33}$ |
| Coverage A | −0.976 | 0 | −0.966 | 0 |
| Metric B | 0.722 | 0 | 0.718 | 0 |
| CAPG | | | 0.923 | 0 |
| **Heterozygous B** | | | | |
| Identifiable | 44 775 | 0 | 127 | $1.9 \times 10^{-29}$ |
| Coverage B | −0.975 | 0 | −0.966 | 0 |
| CAPG | | | 0.944 | 0 |
| **Allelic A** | | | | |
| Hand-Verified | 0.771 | $7.6 \times 10^{-6}$ | 0.685 | $6.9 \times 10^{-6}$ |
| Identifiable | 9.22 | 0.0024 | 0.690 | 0.41 |
| Coverage A | −0.073 | 0 | −0.0480 | 0 |
| Mismatch | 0.0512 | 0 | 0.0113 | $1.2 \times 10^{-89}$ |
| Metric B | 0.0470 | 0 | 0.0332 | 0 |
| CAPG | | | 0.355 | 0 |
| **Allelic B** | | | | |
| Identifiable | 7.12 | 0.0076 | 0.626 | 0.43 |
| Coverage B | −0.069 | 0 | −0.0537 | 0 |
| Mismatch | 0.0510 | 0 | 0.008 57 | $2.6 \times 10^{-54}$ |
| CAPG | | | 0.368 | 0 |
| **Homoeologous** | | | | |
| Hand-Verified | 0.828 | $4.4 \times 10^{-6}$ | 0.695 | $1.2 \times 10^{-4}$ |
| Identifiable | 72.7 | $1.5 \times 10^{-17}$ | 85.1 | $2.8 \times 10^{-20}$ |
| Coverage | −0.485 | 0 | −0.796 | 0 |
| Mismatch | 0.228 | 0 | 0.231 | 0 |
| CAPG | | | 0.379 | 0 |

Table S5: Statistical tests of metric associations for peanut data. Unless otherwise specified, the statistic reported is the Spearman correlation and the $p$-value is for the corresponding hypothesis of no correlation. For association with hand-verified loci and presence of a subgenomic reference mismatch, the $p$-value is from the nonparametric Wilcox rank sum test. For association with identifiable loci, the statistic and $p$-value of a Levene's test of equal variance is reported. We examine heterozygous metrics Eq. 4 from A and B subgenomes, allelic SNP metrics Eq. (5) from A and B subgenomes, and homoeologous SNP metric Eq. (6). Row Hand-Verified reports association between metric and manually assessed truth, as marked in Figure 5. We only hand-verified heterozygous or SNP loci within subgenome A and could confirm no true heterozygous calls. Row Identifiable reports association between the metric and the identifiability of the locus, *i.e.*, whether there is a homoeologous locus within read distance. Row Coverage A/B reports association between the metric and some measure of coverage at the locus. Row Mismatch reports association between the metric and the presence of a mismatch between the subgenomic references. Row Metric B reports correlation with the corresponding metric at the homoeologous locus in subgenome B. Row CAPG is for reporting the correlation between GATK and CAPG metrics.

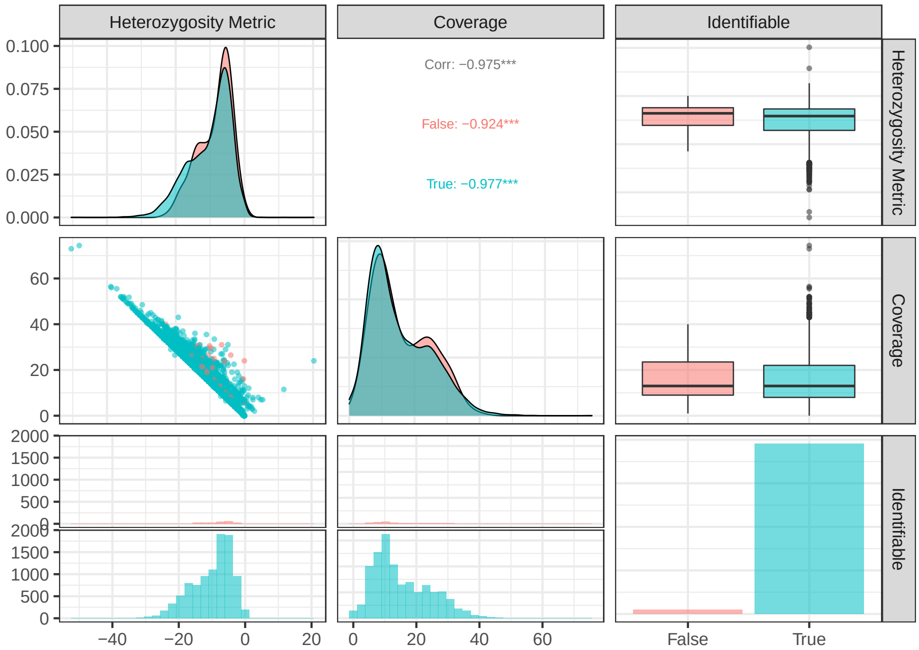

(a) CAPG

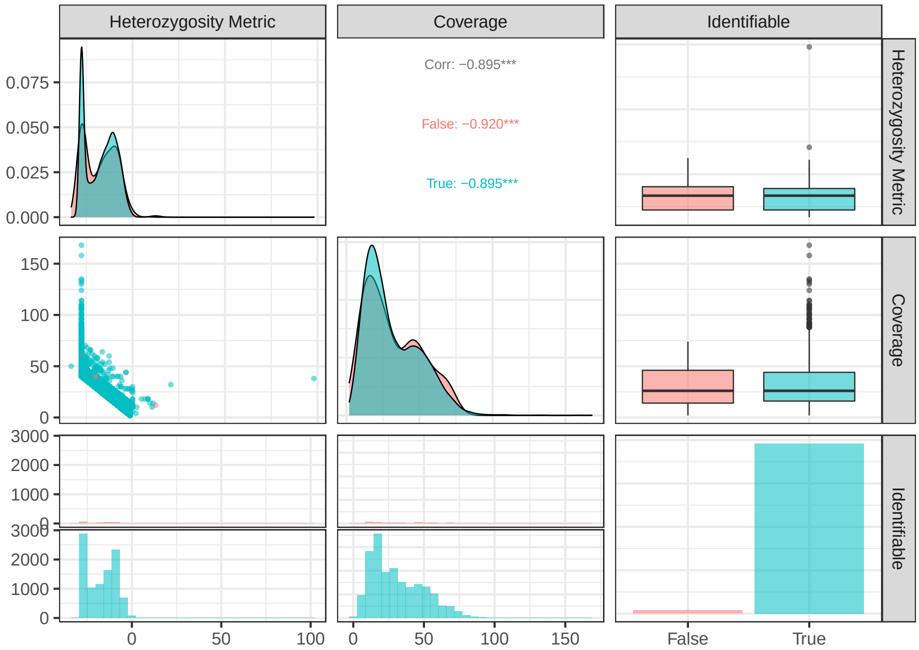

(b) GATK

Figure S7: Pairs plots for peanut of subgenome A (a) CAPG heterozygosity metric Eq. 4 or (b) GATK heterozygosity metric Eq. S3 vs. coverage at locus and an indicator of identifiability, *i.e.*, whether there is a subgenomic difference within read distance. Colored by identifiability status. An unbiased sample of size 10,000 loci is plotted here.

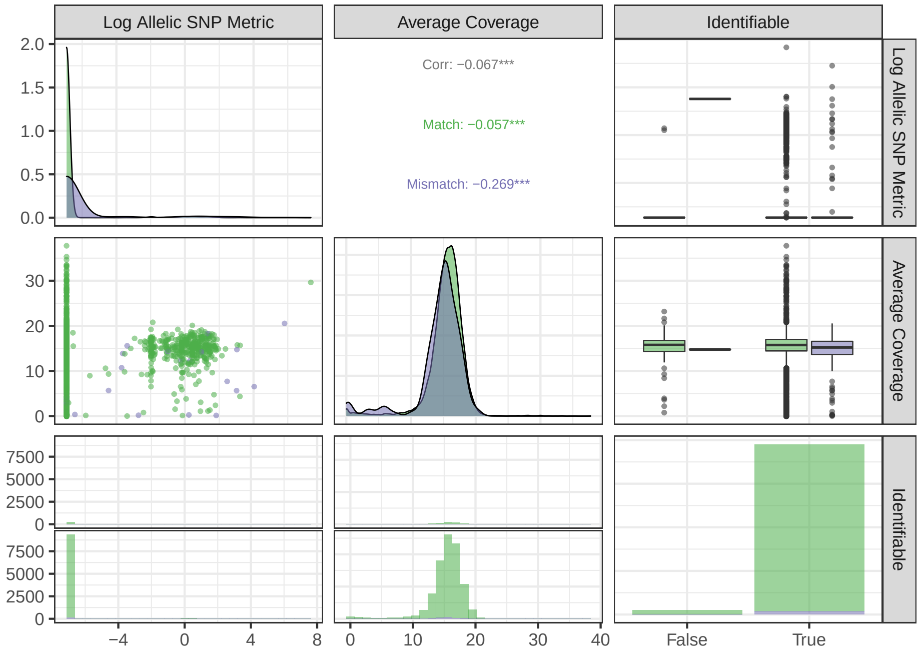

(a) CAPG

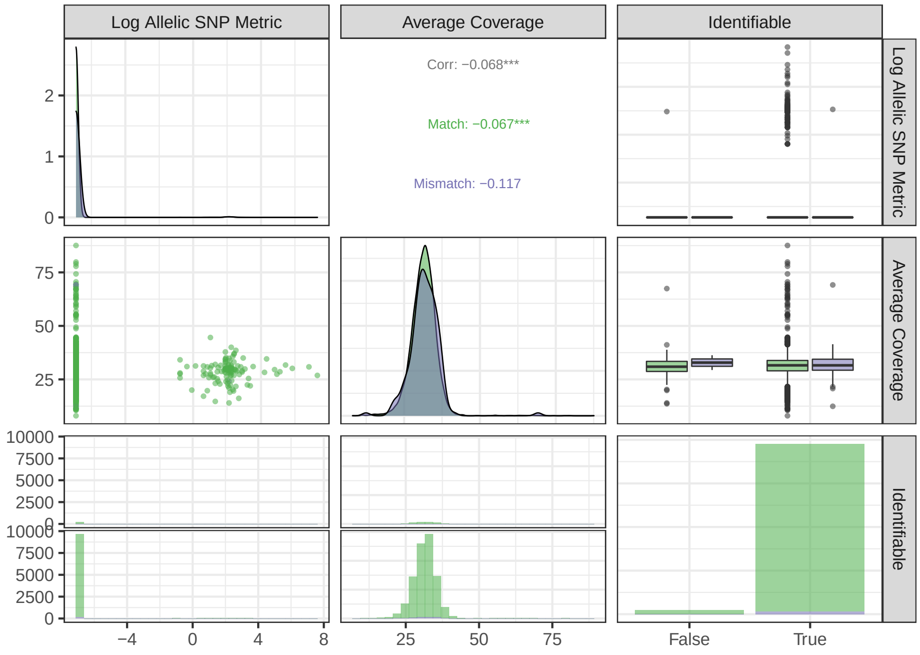

(b) GATK

Figure S8: Pairs plots for peanut of subgenome A (a) CAPG allelic metric Eq. 5 or (b) GATK allelic metric Eq. S4 vs. average coverage and an indicator of identifiability, *i.e.*, whether there is a subgenomic difference within read distance, in subgenome A. Coloring indicates whether the locus is a match (green) or mismatch (purple) between the subgenomic references, which is a fuzzy indicator of truth since mismatches are possible SNPs. An unbiased sample of size 10,000 loci is plotted here.

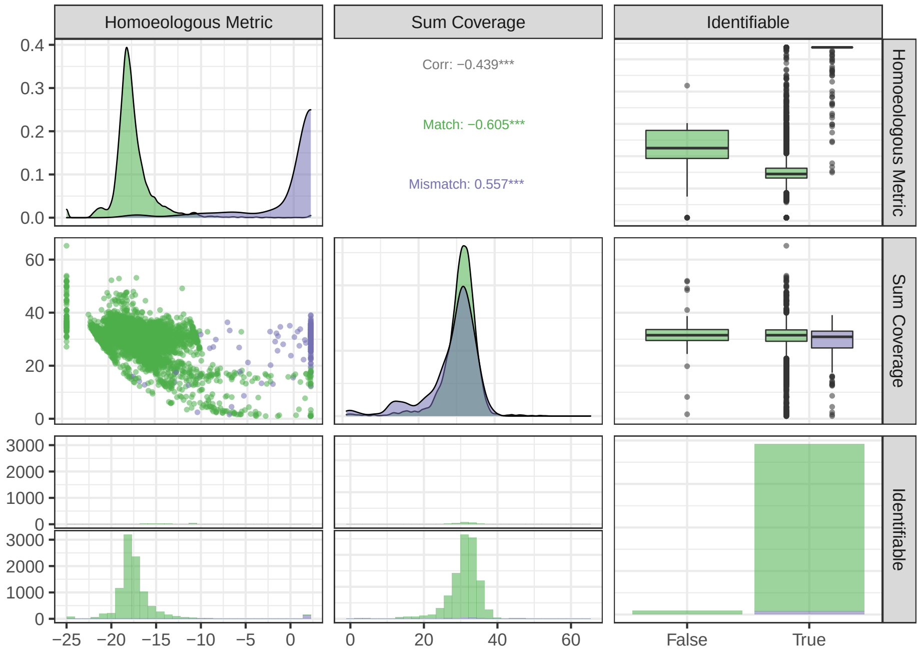

(a) CAPG

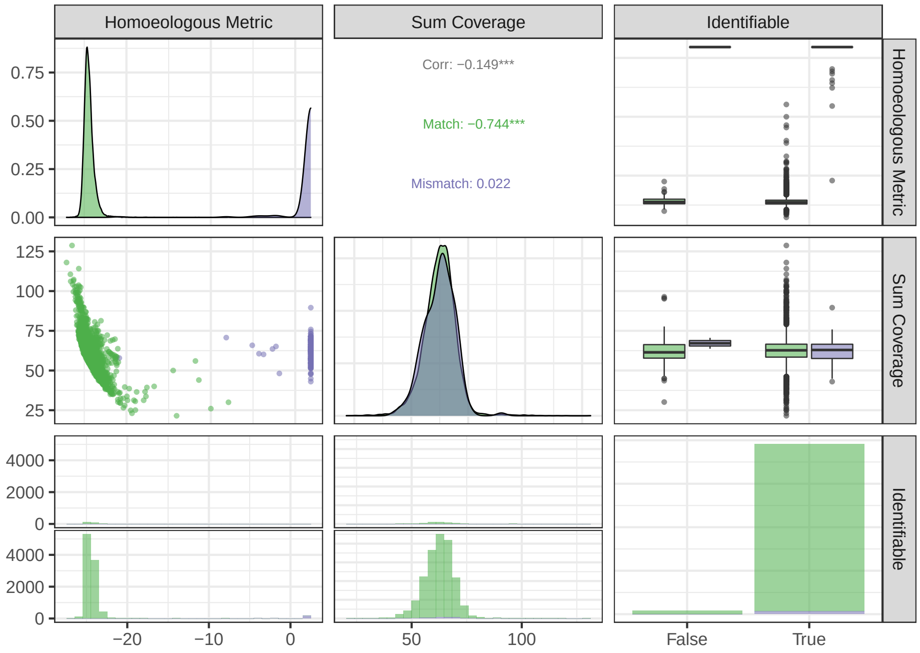

(b) GATK

Figure S9: Pairs plots for peanut of (a) CAPG homoeologous metric Eq. 6 or (b) GATK homoeologous metric Eq. S5 vs. summed average coverage across subgenomes and an indicator of identifiability, *i.e.*, whether there is a subgenomic difference within read distance. Coloring indicates whether the locus is a match (green) or mismatch (purple) between the subgenomic references, which is a fuzzy indicator of truth since mismatches are possible SNPs. An unbiased sample of size 10,000 loci is plotted here.

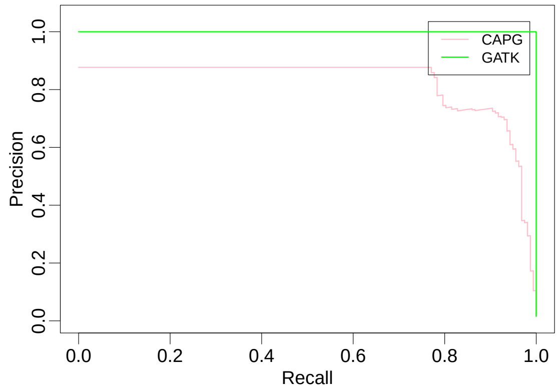

(a) Homoeologous SNPs

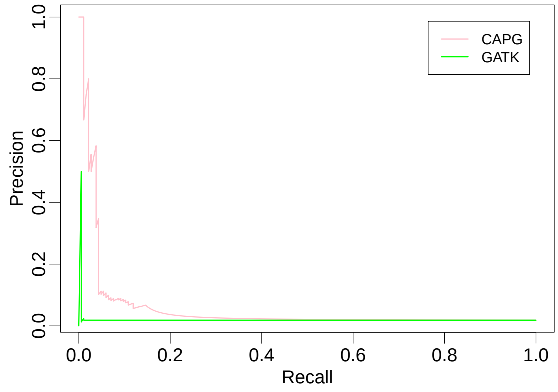

(b) Allelic SNPs

Figure S10: Precision-Recall (PR) curves for predicting (a) homoeologous or (b) allelic SNPs with reference mismatch indicating "true" homoeologous or "true" allelic SNP in peanut. Pink is CAPG, green is GATK.

H. Levene. Robust tests for equality of variances. In I. Olkin, editor, *Contributions to Probability and Statistics*, pages 278–292. Stanford University Press, Palo Alto, CA, 1960.

H. Li, B. Handsaker, A. Wysoker, T. Fennell, J. Ruan, N. Homer, G. Marth, G. Abecasis, and R. Durbin. The sequence alignment/map format and SAMtools. *Bioinformatics*, 25(16):2078–2079, 2009.

P. Lindenbaum. JVarkit: Java-based utilities for bioinformatics, 2015.

G. Marçais, A. L. Delcher, A. M. Phillippy, R. Coston, S. L. Salzberg, and A. Zimin. MUMmer4: A fast and versatile genome alignment system. *PLoS Computational Biology*, 14(1):e1005944, 2018.

R. Nielsen, J. S. Paul, A. Albrechtsen, and Y. S. Song. Genotype and SNP calling from next-generation sequencing data. *Nature Reviews Genetics*, 12(6):443–451, 2011.
